# H3K27me3 dictates atypical genome-nuclear lamina interactions and allelic asymmetry during early embryogenesis

**DOI:** 10.1101/2023.02.06.527307

**Authors:** Isabel Guerreiro, Franka J. Rang, Yumiko K. Kawamura, Femke C. Groenveld, Ramada E. van Beek, Silke J. A. Lochs, Ellen Boele, Antoine H. M. F. Peters, Jop Kind

**Author notes:** These authors contributed equally (Isabel Guerreiro, Franka J. Rang).

## Abstract

The very first days of mammalian embryonic development are accompanied by epigenetic reprogramming and extensive changes in nuclear organization. In particular, genomic regions located at the periphery of the nucleus, termed lamina-associated domains (LADs), undergo major rearrangements after fertilization. However, the role of LADs in regulating gene expression as well as the interplay with various chromatin marks during preimplantation development remains elusive. In this study, we obtained single-cell LAD profiles coupled with the corresponding gene expression readout throughout the first days of mouse development. We detect extensive cell-cell LAD variability at the 2-cell stage, which surprisingly does not seem to functionally affect gene expression. This suggests an unusual uncoupling between 3D-nuclear genome organization and gene expression during totipotent developmental stages. By analyzing LAD dynamics and chromatin states across early developmental stages in an allelic-specific manner, we identify genomic regions that transiently detach from the nuclear lamina and are enriched by non-canonical H3K27me3. Upon maternal knock-out of a component of the Polycomb repressive complex 2 and concomitant loss of H3K27me3 during early embryogenesis, these regions relocate to the lamina at the 2-cell stage. Our results suggest that H3K27me3 is the prime determinant in establishing the atypical distribution of the genome at the nuclear periphery during the first days of embryonic development. This study provides insight into the molecular mechanisms regulating nuclear organization of parental genomes during very early mammalian development.

## Main

Mammalian development begins with the fusion of two differentiated cells, the gametes, that give rise to a totipotent zygote. The embryo subsequently undergoes multiple cycles of cell division, with inner cells progressively transitioning to a pluripotent state while outer cells commit to becoming extra-embryonic tissue by the time of implantation in the uterus. After fertilization, maternal transcripts are progressively degraded as embryonic genes become active. All these events occur within the first three days of mouse development and are accompanied by extensive epigenetic reprogramming as well as major changes in spatial genome organization (reviewed in ^1, 2^).

One important feature of nuclear organization is the localization of genomic regions at the nuclear lamina (NL). These genomic regions, termed lamina-associated domains (LADs), have been extensively studied and are characterized by being gene-poor, having low gene expression and high repeat content as well as other features of constitutive heterochromatin. Importantly, in addition to their architectural function, LADs have been shown to play a role in gene expression (reviewed in ^3–5^). LADs are typically detected using the DamID technique^6^. This technique allows for the profiling of protein-DNA interactions by fusing a protein of interest to the Dam DNA methyltransferase, a DNA methyltransferase that methylates adenines in a GATC context of genomic regions it comes into contact with. Fusing Dam to a component of the nuclear lamina (NL), typically Lamin B1, thus results in the specific methylation of LADs that can subsequently be sequenced and mapped to the genome.

Previous work studying LADs in the context of preimplantation development has shown atypical lamina association patterns at these stages. Maternal LADs have been found to be established *de novo* following fertilization, while paternal LADs undergo massive rearrangements between the zygote and 2-cell stages. Consequently, maternal and paternal genomes show differences in lamina association up until the 8-cell stage^1^.

The positioning of the genome at the NL or other locations within nuclear space is non-random and is associated with specific chromatin and transcriptional states^7^. Therefore, it is essential to study LADs within the wider context of epigenetics and gene expression. Profiling histone post-translational modifications (PTMs) in early embryogenesis is challenging and recent work has started to shed light on the epigenetic features and dynamics of the early mouse embryo^8–14^. Trimethylation at histone 3 lysine 27 (H3K27me3), a histone modification that is associated with the repression of developmental genes, is deposited by the Polycomb repressive complex (PRC) 2. After fertilization, H3K27me3 has been shown to lose its typical distribution at promoters of developmental genes at maternal and paternal genomes while retaining non-canonical broad distal domains along the maternal genome at regions devoid of developmental genes^14^. Trimethylation at histone 3 lysine 9 (H3K9me3), a mark often found in LADs, also shows unusual enrichment and allelic asymmetry in early mouse development. ChIP-seq data has shown that H3K9me3 and H3K27me3 extensively overlap during early developmental stages across the genome, in contrast to what has been reported in other biological systems^12^. Although low-input technologies have recently shed light on the chromatin state and nuclear architecture of the early mouse embryo, the underlying mechanisms and the relationship between the different layers of epigenetic features remains largely unexplored. Additionally, while cell-cell variability in gene expression and chromatin modifiers is proposed to contribute to early cell fate choices^15–17^, the role of variable genome-NL interactions during preimplantation development remains largely unexplored. Here, we profile single-cell LADs throughout a range of developmental stages in early development and study their cell-cell variability and dynamics over time in the wider context of chromatin state and transcription.

### Genome-nuclear lamina association is highly variable among single cells of 2-cell embryos

Our previous work profiling LADs in preimplantation development has suggested that cell-cell variability in LADs may be particularly high between single cells in the early developmental stages^1^. To address this finding in more detail we have made use of scDam&T-seq, a newly developed single-cell DamID technique^18^ which: 1) provides improved signal-noise ratio, 2) increases the throughput, 3) allows to keep track of the embryo each cell comes from and 4) can be coupled with the corresponding transcriptional output from the same cell (Extended Data Figure 1a-b). Using this technique we obtained a total of 754 single-cell LAD profiles that passed quality control thresholds (see Methods): 107 zygote cells, 196 2-cell stage cells, 183 8-cell stage cells and 268 mES cells (Extended Data Figure 1b-c). Average LAD profiles per stage showed high concordance with previously published data (Extended Data Figure 1e-f). For each collected cell we obtained the LAD profile and the corresponding gene expression read-out (Figure 1a and Extended Data Figure 2a). Plotting all single cells by uniform manifold approximation and projection (UMAP) based on DamID (i.e. Dam-LMNB1 value) shows clear clustering according to stage (Figure 1b), similarly to the UMAP visualization obtained from the gene expression information of the same cells (Figure 1c). Thus, consistent stage-specific LAD patterns are observed between individual cells of the same cleavage stage. The percentage of the genome that contacts the nuclear lamina was similar across stages and comparable to mouse embryonic stem cells (mESCs) with the exception of the zygote which had a markedly lower proportion of the genome located at the NL (Extended Data Figure 2b).

**Figure 1.**
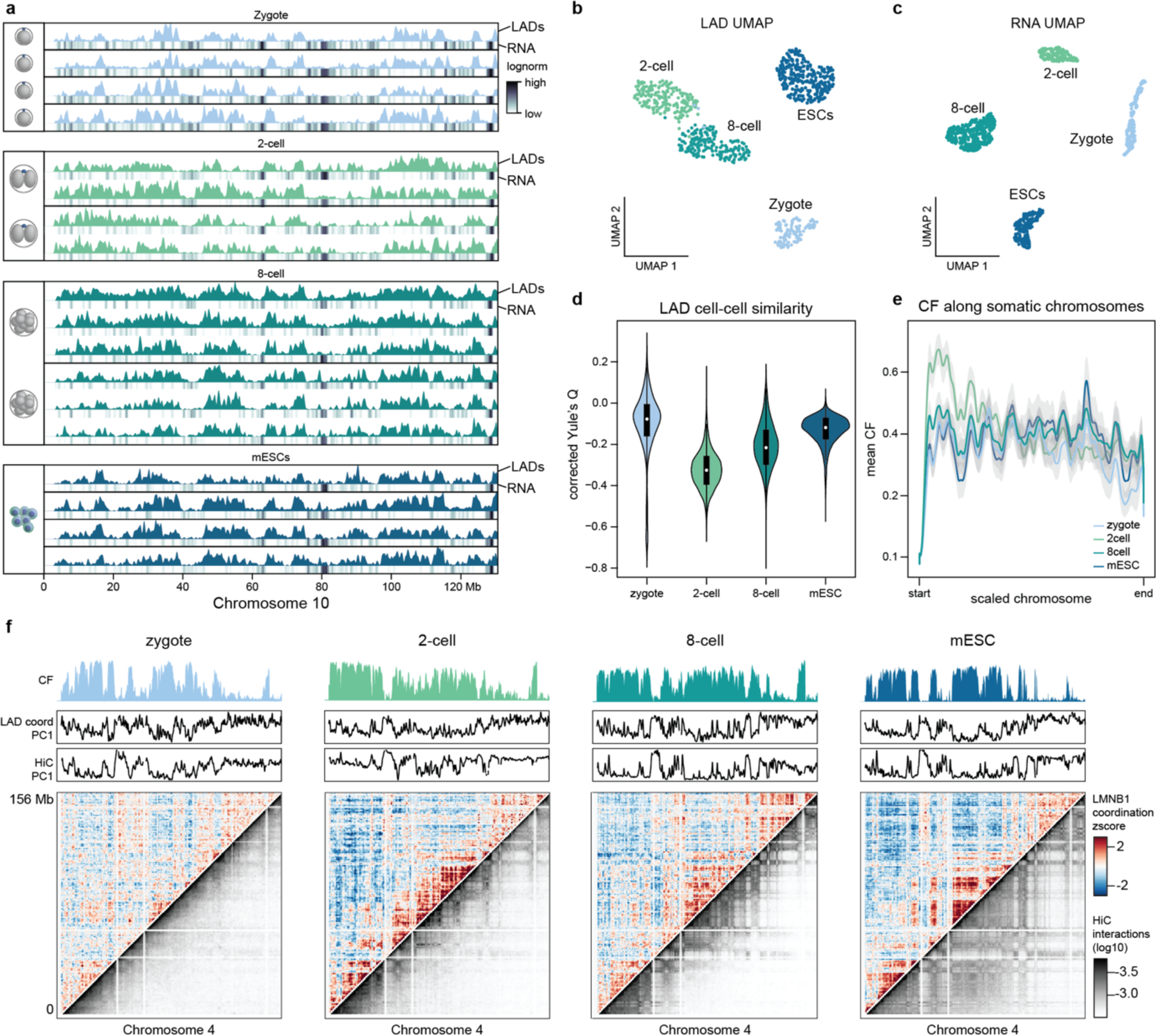
Genome-nuclear lamina contacts at the two-cell stage are very variable among single cells. **a,** Examples of LAD single-cell profiles (RPKM) and corresponding gene expression track (log-normalized RPKM scaled to maximum value per sample) across the entire chromosome 10 at different developmental stages and mESCs. **b,** UMAP based on Dam-LMNB1 single cell readout. **c,** Single-cell UMAP based on transcriptional readout of the same cells as (b). **d,** Distribution of corrected Yule’s Q of the Dam-LMNB1 single-cell data providing a measure for LAD cell-cell similarity per stage. **e,** Smoothed mean (1000-Mb Gaussian kernel) of LMNB1 contact frequency (CF) distribution across all autosomal chromosomes scaled to the same size per stage. **f,** comparison between LMNB1 coordination (red, white, blue scale) and Hi-C interaction (grey scale) matrices along the entire chromosome 4 for all stages. Principal component 1 for each of the metrics is shown above as well as LMNB1 CF.

To understand whether there were differences in cell-cell LAD variability across stages, we converted single-cell LMNB1 values to an aggregate contact frequency (CF) which refers to the proportion of cells for which a genomic bin is contacting the NL^19^.

CF distributions per stage revealed more intermediate CF values for 2-cell and 8-cell stages, indicating more variability in lamina association (Extended Data Figure 2c). To more precisely quantify LAD variability, we calculated the Yule’s Q coefficient which provides a measure of similarity between single-cell LADs. To control for the different levels of sparsity and noise per stage, we corrected Yule’s Q values using the same metric on randomised single-cell profiles (Methods and Extended Data Figure 2h). This approach allows us to discern true single-cell variability from noise-driven variability and showed that 2-cell LADs are particularly heterogeneous among single cells compared to other stages (Figure 1d and Extended Data Figure 2d). Strikingly, we also found that cells from the same embryo tend to have more similar LAD profiles, especially at the 2-cell stage (Extended Data Figure 2e). This could potentially be due to restraints dictated by nuclear organization and chromatin states present in the zygote prior to cell division. However, we cannot completely exclude a technical component since the volume of injected Dam construct per embryo may slightly vary.

Lastly, to determine whether the level of variability was constant throughout the linear chromosome, we plotted the distribution of CFs across all autosomal chromosomes at the 2-cell stage and found that genomic regions proximal to the centromere showed unusually high CF values (Figure 1e and Extended Data Figure 2f). Similarly, contacts with the nuclear lamina in the first 30 Mb (centromeric) were significantly higher at the 2-cell stage compared to CF values along the remaining portion of the chromosome (non-centromeric), unlike mESCs which showed no significant difference in nuclear lamina-contacts between centromeric and non-centromeric regions (Extended Data Figure 2g). These results suggest that despite high levels of LAD variability, centromeric regions tend to associate with the lamina in a more uniform manner across cells. This is a feature uniquely observed at the 2-cell stage, which coincides with the moment in development where centromeric regions dramatically change their organization in the nucleus, beginning to relocate from the border of nucleolar precursor bodies to cluster into chromocenters^20^.

Having found a pattern in the LAD variability along the chromosome we next asked whether LADs would vary independently of each other. Previous work on a human cell line has shown that genomic regions that are in close spatial proximity coordinately attach and detach from the NL in single cells. As a result, LAD coordination measurements show a similar pattern to genome interactions visualized by Hi-C DNA-DNA matrices^19^. After fertilization, chromatin is mostly unorganized and progressively acquires higher-order chromatin structure as development progresses^21^. Interestingly, LADs were shown to be established already in the zygote as big domains of high A/T content, thus preceding the establishment of 3D-genome topology^1^. This fact prompted us to ask what the interrelationship is between 3D-genome topology and LAD coordination at these early stages of development. By calculating the coordination between lamina-association values per single cell, we found that although LADs are clearly present at the zygote stage, the coordination values are very low and tend to increase over developmental time (Figure 1f). This suggests that LAD coordination and 3D-genome organization are interconnected. Indeed, when we performed a principal component (PC) analysis for both measurements, PC1 of the Hi-C showed high correlation with either PC1 or PC2 of the LAD coordination matrix (Figure 1f and Extended Data Figure 2j).

Previously, we also found that genome-NL contacts interact multivalently over long stretches in single cells of a human cell line^19^. We thus wondered if this would also be the case during preimplantation development, in the near absence of higher-order chromatin organization. We found that, at all stages, longer stretches of the genome are in contact with the nuclear lamina when compared to a randomized control, including the zygotic stage (Extended Data Figure 2i). This shows that unlike LAD coordination, multivalent genome-NL contacts appear to be established largely independent of 3D-genome topology.

These results indicate that LADs may be established through multivalent interactions just after fertilization independent of higher order chromatin organization, but that the mechanisms driving 3D-genome topology are tightly interconnected with LAD coordination from the 2-cell stage onwards.

### Cell-cell LAD variability at the 2-cell stage is higher in the paternal allele

Previous work has reported parental differences in genome-lamina association up to the 8-cell stage^1^. This prompted us to investigate single-cell LAD variability in the maternal and paternal alleles by using a hybrid cross between mice of two different strains (CBAxC57BL/6J females and CAST/EiJ males). For mESCs a hybrid strain of Cast/EiJx129Sv was used. We could confirm the previously reported LAD differences between alleles, which are apparent even at the single-cell level (Figure 2a and Extended Data Figure 3a and b). Visualization of all single-cell allele-specific LAD profiles by UMAP showed a clear allelic separation at the zygote and 2-cell stages in contrast to the intermingling of the paternal and maternal LADs in ESCs (Extended Data Figure 3d). Paternal and maternal LADs at the zygote stage clustered separately from each other. Interestingly, paternal LADs grouped closer to mESC LADs implying that paternal genome displays more canonical lamina association at the zygote stage. After cell division, at the 2-cell stage, paternal genome-NL interactions are largely rearranged^1^. Both in zygote and 2-cell stages, the paternal genome contacts the NL significantly more than the maternal counterpart, while in ESCs LAD coverage was comparable between the two alleles (Figure 2b). Interestingly, while at the zygote stage maternal LADs are more variable than paternal LADs, this trend is clearly inverted at the 2-cell stage (Figure 2c).

**Figure 2.**
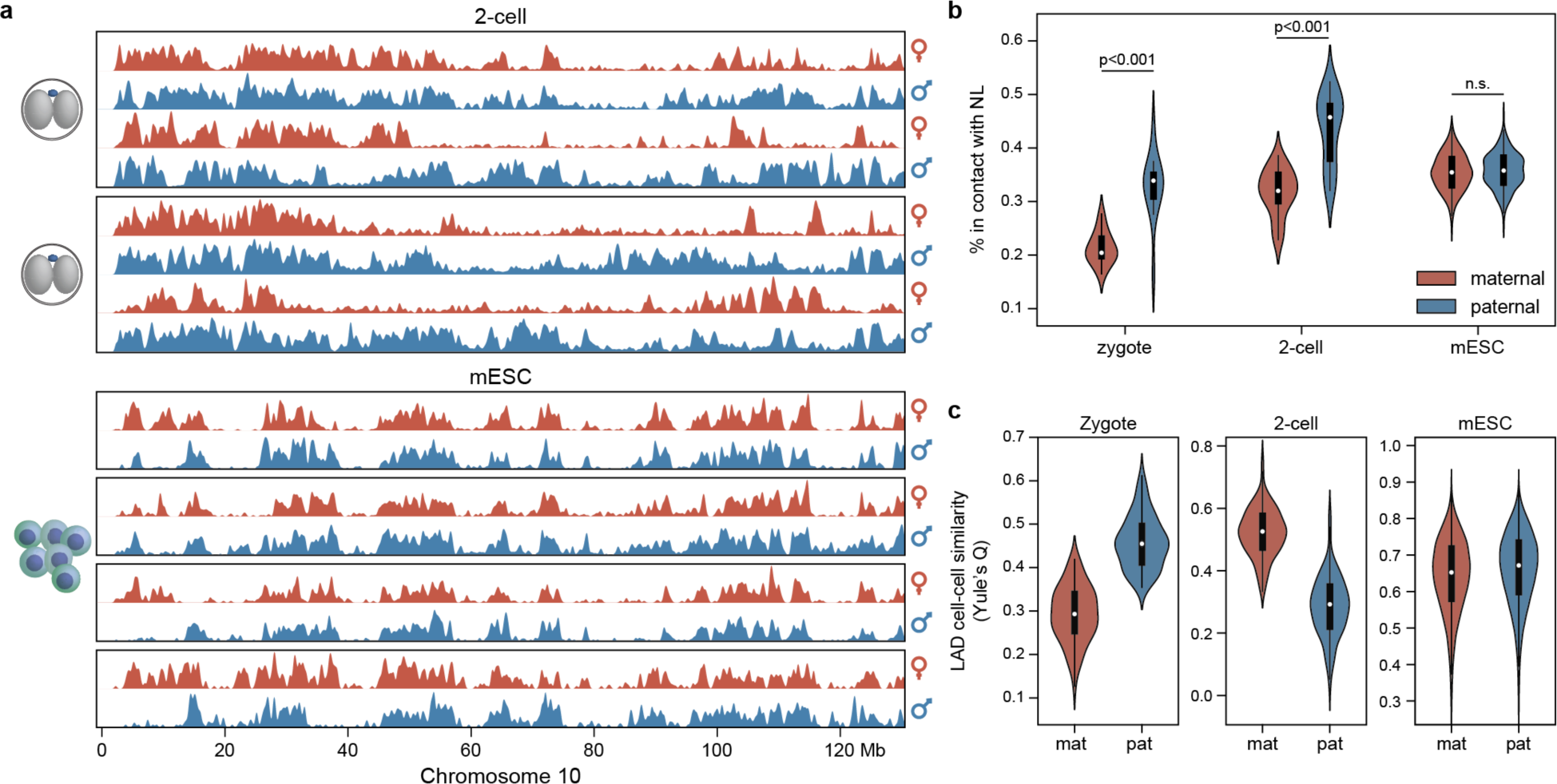
Paternal LADs are more variable across single cells than maternal LADs at the 2-cell stage. **a,** 2-cell and mESC example single-cell LAD profiles split in maternal (red) and paternal (blue)alleles. For the two-cell stage, allele-specific profiles from the same embryo (two cells) are enclosed in a blackbox while for mESCs allele specific profiles of each single cell are enclosed in a black-box. **b,** percentage of the genome that is in contact with the NL per allele and per stage. Two-sided Wilcoxon rank-sum test was performed (zygote stage, n=14, p-value=0.00014; 2-cell stage, n=26, p-value=1.83×10^-07^; mESC, n=268, p-value=0.33). **c,** Violin plots with distribution of Yule’s Q values as a measure of LAD cell-cell similarity for the maternal (red) and paternal (blue) alleles. Two-sided Wilcoxon rank-sum test was performed (zygote stage, n=14; 2-cell stage, n=26; mESC, n=268) (Extended Data Table 6).

Due to the asymmetry in LAD content and variability between the two parental alleles, we wondered whether the tendency for high CF values at centromeric regions of chromosomes at the 2-cell stage was a feature of both alleles. Indeed, the maternal as well as the paternal Dam-LMNB1 CF values were higher in centromeric regions of chromosomes at the 2-cell stage compared to zygote and mESCs (Extended Data Figure 3e). However, the difference in CF between centromeric and non-centromeric regions was more noticeable in the maternal genome (Extended Data Figure 3f), presumably as a result of the paternal genome having generally higher NL association. Interestingly, even though the maternal zygotic LAD profile has been shown to resemble that of the 2-cell stage^1^, centromere-specific lamina associations were not present at this stage.

Overall, these results indicate that the parental genomes do not locate evenly to the nuclear periphery and that paternal LADs contribute the most to the unusually high cell-cell variability observed at the 2-cell stage (Figure 1d).

### LAD variability at the 2-cell stage is not accompanied by major changes in chromatin state and transcription

The localization of genomic regions at the NL is typically associated with heterochromatic features and low gene expression. Since LAD cell-cell variability is unusually high at the 2-cell stage, we hypothesized that differential NL association may impact chromatin state or gene expression. For this, we employed EpiDamID^22^, a single-cell DamID-based technique that has been adapted to detect histone marks through the fusion of either single-chain variable fragments (scFv) or chromatin reader domains to the Dam methylase. We chose to profile 1) H3K9me3 that is often present in LADs, 2) H3K27me3 which plays an essential role in repressing genes during embryonic development and 3) open chromatin that tends to anti-correlate with LADs. To this end, the fusion constructs Dam-Cbx1 (chromodomain tuple) and Dam-H3K27me3 (scFv)^22^ as well as the untethered Dam were used, respectively. The profiles obtained with these constructs were comparable to published 2-cell stage ChIP-seq and ATAC-seq data (Figure 3a and Extended Data Figure 4a). Importantly, a mutant version of the Dam-H3K27me3 construct^22^, which loses binding affinity to the histone mark, showed no specific enrichment in contrast to the non-mutant construct (Extended Data Figure 4c). The DamID datasets had high genome-wide correlations with the corresponding publicly available dataset, which further validated their genome-wide similarity (Figure 3b).

**Figure 3.**
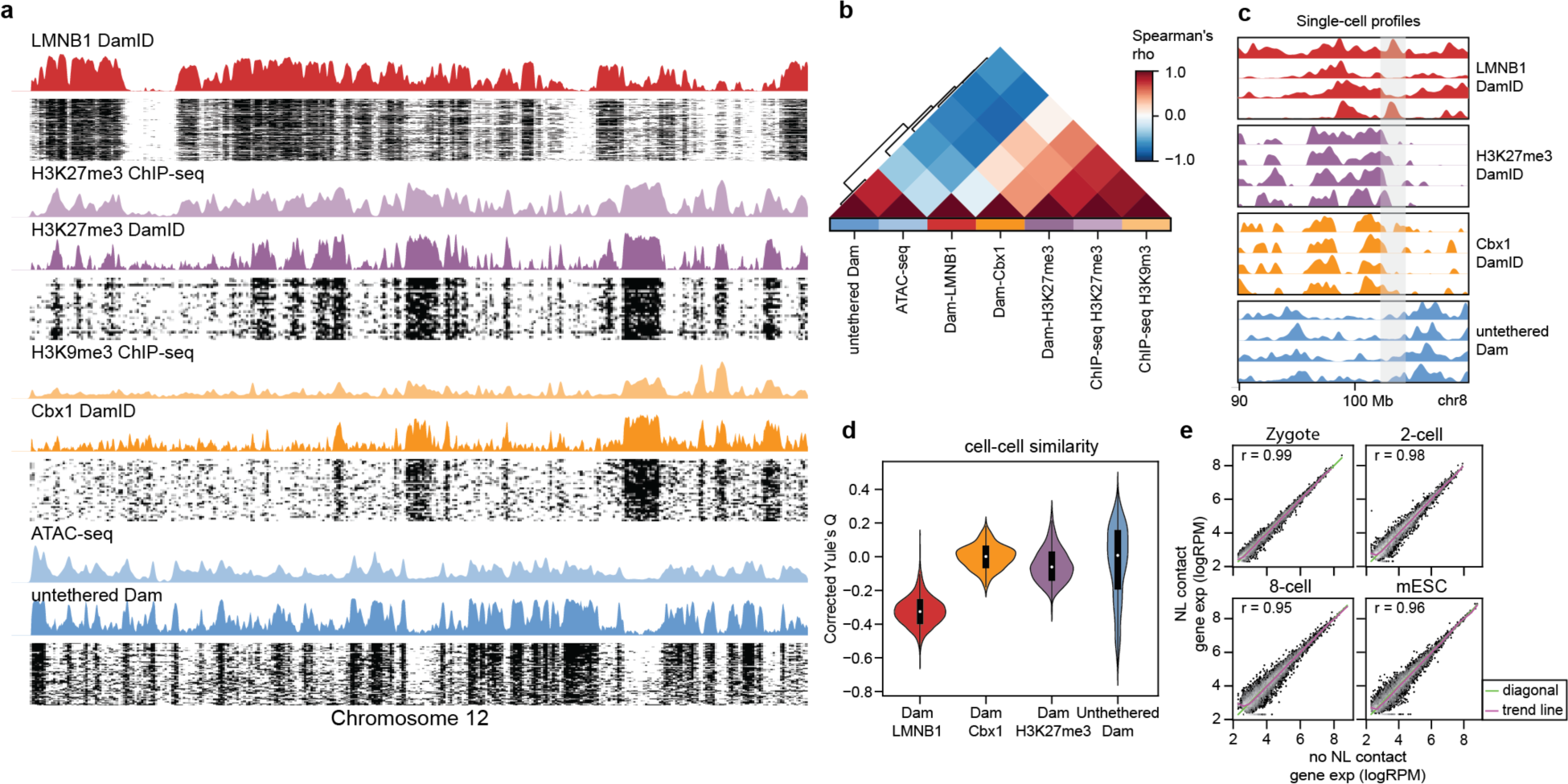
LAD variability in single-cells does not result in chromatin state or gene expression changes at the 2-cell stage. **a,** Heatmap of binarized single-cell profiles for Dam-LMNB1, Dam-H3K27me3, Dam-Cbx1 across the entire chromosome 12 ordered by decreasing unique number of GATCs. All cells that passed quality control thresholds are plotted and corresponding CF values are shown above each heatmap. When available, the corresponding ChIP-seq publicly available dataset (RPKM) was plotted on top of the DamID track. All data originates from the 2-cell stage. **b,** Spearman correlation heatmap relating DamID (present study) and corresponding ChIP-seq (previous studies) measurements at the 2-cell stage. **c,** Genome browser view of four example single cells per construct over a selected region in chromosome 8. A particularly variable region is highlighted with a grey shading. **d,** Distribution of corrected Yule’s Q of the single-cell data providing a measure for cell-cell similarity per construct. **e,** Gene expression comparison between cells that have a genomic region at the NL and cells for which that same genomic region does not contact the NL. Correlation was calculated using Pearson’s r. The trend line of the data is in purple and the diagonal expected for a one-to-one correspondence of the data is in green.

Strikingly, when inspecting highly variable regions with respect to genome-NL interactions, H3K27me3, H3K9me3 and open chromatin appeared to have a more uniform distribution among single cells (Figure 3c). In order to quantify and compare the levels of cell-cell variability among the different measurements, we again controlled for construct-specific sparsity and noise by normalizing the Yule’s Q of our data to the same metric from a randomized dataset (Extended Data Figure 4a and Methods). Visual comparison of the single-cell LAD profiles with the corresponding randomized data revealed clear differences that were less evident for the other constructs (Extended Data Figure 4a). The normalized cell-cell similarity measurement confirmed these observations as LADs showed a lower similarity score compared to H3K9me3, H3K27me3 and accessible chromatin (Figure 3d and Extended Data Figure 4d). These results indicate that LAD variability among single cells at the 2-cell stage is not accompanied by changes in chromatin state.

We hypothesized that variable lamina association could result in transcriptional differences among single cells. To test this, we made use of the combined genomic and transcriptomic read-out of our single-cell data (Figure 1a). We validated our gene expression data by calculating enrichment of genes known to be activated at different timepoints along the first days of development. The results showed stage-specific upregulation of the different categories of genes at the correct developmental time (Extended Data Figure 4e). However, when comparing the gene expression of genomic regions when they reside at the lamina versus when they are dissociated, we observe no noticeable differences (Figure 3e).

Together these results indicate that while genome-NL associations show high cell-cell variability at the 2-cell stage, this does not result in large changes in chromatin state or gene expression.

### Regions that dissociate from the NL at the 2-cell stage are high in H3K27me3

LADs in somatic cells typically display features of constitutive heterochromatin^23^. Interestingly, most chromatin marks show an atypical distribution during the first days of embryonic development^9, 10, 12–14^. This fact motivated us to investigate the relationship between LADs and histone PTMs during preimplantation development. To that end we performed clustering of genomic bins based on their Dam-LMNB1 and histone PTM values across stages (Methods). We could assign 8 clusters (Extended Data Figure 5a-c) that are characterized by different levels of lamina association and histone PTMs. We focused on the four clusters that showed high Dam-LMNB1 levels at least in one of the stages (Figure 4a). Clusters 1 and 2 include LADs that can be observed in all stages - constitutive LADs – and show mild enrichment for maternal H3K9me3 (cLADs-K9) and H3K4me3 (cLADs-K4), respectively (Figure 4a, Extended Data Figure 5c). Cluster 3 contains genomic regions that preferentially associate to the NL at the 2-cell stage and to a lesser degree at the 8-cell stage, which we termed embryonic transient LADs or ET-LADs. ET-LADs were the only LAD clusters to show low LINE L1 density and higher gene density, contrary to the features that typically characterise LADs (Extended Data Figure 5e-f). In addition, ET-LADs appear to be enriched in H3K27me3 in ESCs, where these regions have mostly lost lamina association (Extended Data Figure 5b). Cluster 4 represented genomic regions in the maternal genome that are associated to the NL in mESCs but not during early developmental stages, namely zygote and 2-cell stage (Figure 4b and Extended Data Figure 5c, upper panel). We therefore termed these regions embryonic transient iLADs or ET-iLADs. Although these regions are not LADs in the maternal zygotic genome, they do show strong lamina association in the paternal zygotic genome (Extended figure 5c, bottom panel). At the 8-cell stage these regions appear to show mild lamina association values suggesting an intermediate state of NL attachment (Figure 4b). Interestingly, ET-iLADs showed strong H3K27me3 signal at stages with reduced lamina association levels (Figure 4a, c and Extended Data Figure 5b). These genomic regions display an inverse relationship between lamina association and the non-canonical H3K27me3 (ncH3K27me3) that is found in early development up to implantation stages ^14^. Indeed, we found that regions that had high lamina association values in mESCs but not at the 2-cell stage, were instead H3K27me3-enriched at the 2-cell stage (Figure 4d-f). Additionally, we found a depletion of Dam-LMNB1 signal over maternal 2-cell ncH3K27me3 domains at the two-cell stage which was in contrast with the enrichment observed in mESCs and zygote (Extended Data Figure 5d). Altogether these results indicate that a group of genomic regions specifically detach from the NL during the first days of embryonic development while being enriched with non-canonical H3K27me3. These observations point towards a link between NL-association and Polycomb regulation during preimplantation development.

**Figure 4.**
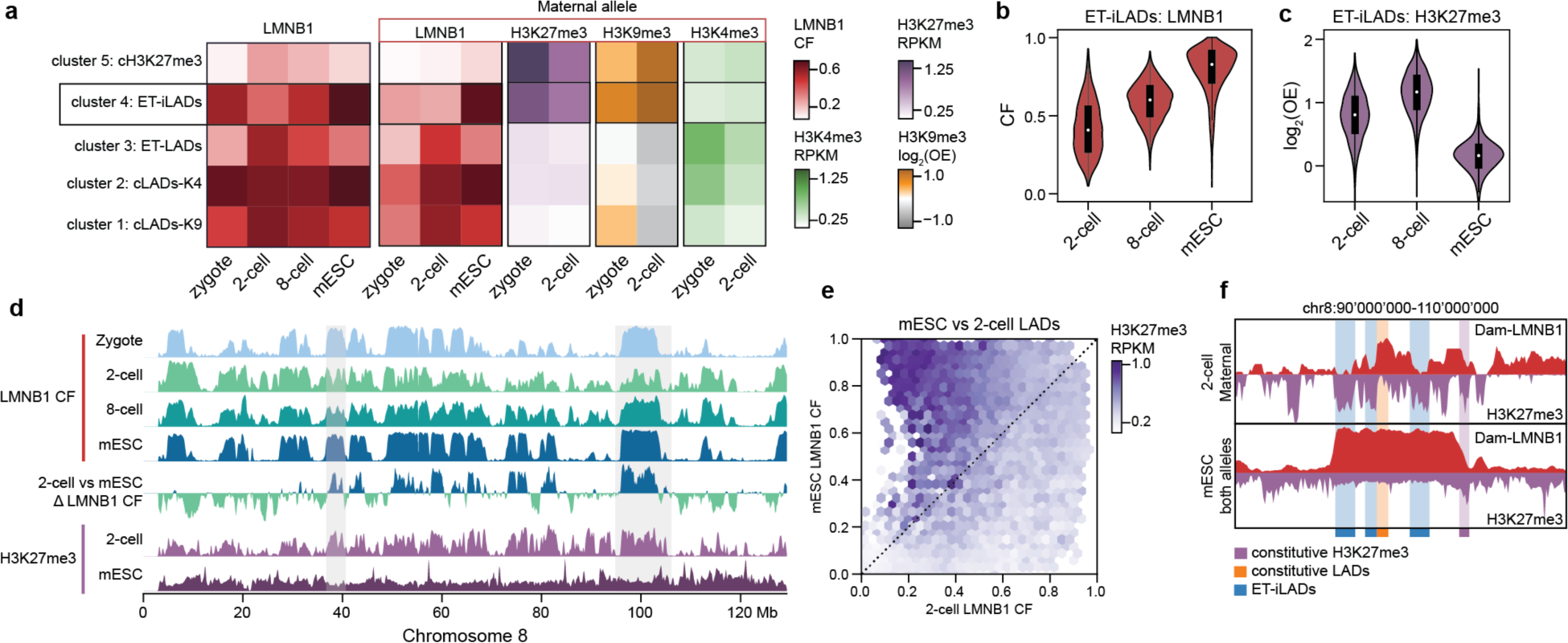
ET-iLADs that specifically detach from the NL in early embryos are enriched in H3K27me3. **a,** Clustering of genomic regions based on their NL attachment and epigenetic features. Average values per identified cluster was calculated for each stage. Five of the eight clusters are depicted in this panel. On the left are the LMNB1 CF values from both alleles combined and on the right maternal LMNB1 CF, H3K27me3, H3K9me3 and H3K4me3 values from the maternal allele are plotted. An extended version of this plot is shown in Extended Data Figures 5b and c. **b,** Violin plot showing the LMNB1 CF values in ET-iLADs per stage. **c,** Violin plot showing the H3K27me3 RPKM values in ET-iLADs per stage. **d,** Dam-LMNB1 CF profiles of all stages and differential CF profile between mESC and 2-cell stage values along chromosome 8. H3K27me3 profiles of the 2-cell stage and mESCs are also plotted. **e,** Genome-wide comparison between mESC and 2-cell LMNB1 CFs. Color intensity refers to H3K27me3 RPKM values. **f,** Example genomic region where three of the genomic clusters identified in (a) can be visualized. Mirror profiles show LAD values on top and H3K27me3 on the bottom for the 2-cell stage (maternal allele) and mESCs (both alleles).

### H3K27me3 sequesters genomic regions away from the NL at the 2-cell stage

Having observed an apparent inverse relationship between NL association and H3K27me3, we proceeded to deplete H3K27me3 in early development using a maternal knockout of *Eed* (*Eed* mKO), an essential component of Polycomb repressive complex 2 (PRC2) that deposits H3K27me3. This mutation is acquired in growing oocytes of *Eed*^fl/fl^;Gdf9^Cre^ female mice and results in H3K27me3 loss from the oocyte stage up to the 8-cell stage^21, 24^. Embryos obtained from crosses with *Eed*^fl/fl^ mothers are used as control. To obtain allelic-resolved data we performed hybrid crosses between C57BL/6J females and JF1/MsJ males.

We performed scDamD&T-seq with the Dam-LMNB1 construct on both *Eed* mKO and control 2-cell embryos to uncover the effect of H3K27me3 absence on LADs. We obtained 95 and 120 single-cell LAD profiles passing quality control thresholds for the mKO and control genotypes (Extended Data Figure 6a). Comparison of LAD profiles between the two conditions showed extensive differences in NL association upon H3K27me3 depletion (Figure 5a). To understand these differences in the context of NL association throughout preimplantation development, we visualized the LAD data together with previously collected samples using UMAP. While *wt* 2-cell maternal and paternal LADs cluster apart, parental LAD profiles from *Eed* mKO embryos cluster closely together as well as with the 2-cell paternal LADs. This suggests that H3K27me3 depletion nearly equalizes allelic differences in LADs. This result is caused by the maternal allele acquiring paternal-like NL association patterns in the absence of H3K27me3 (Figure 5b, and Extended Data Figure 6a).

**Figure 5.**
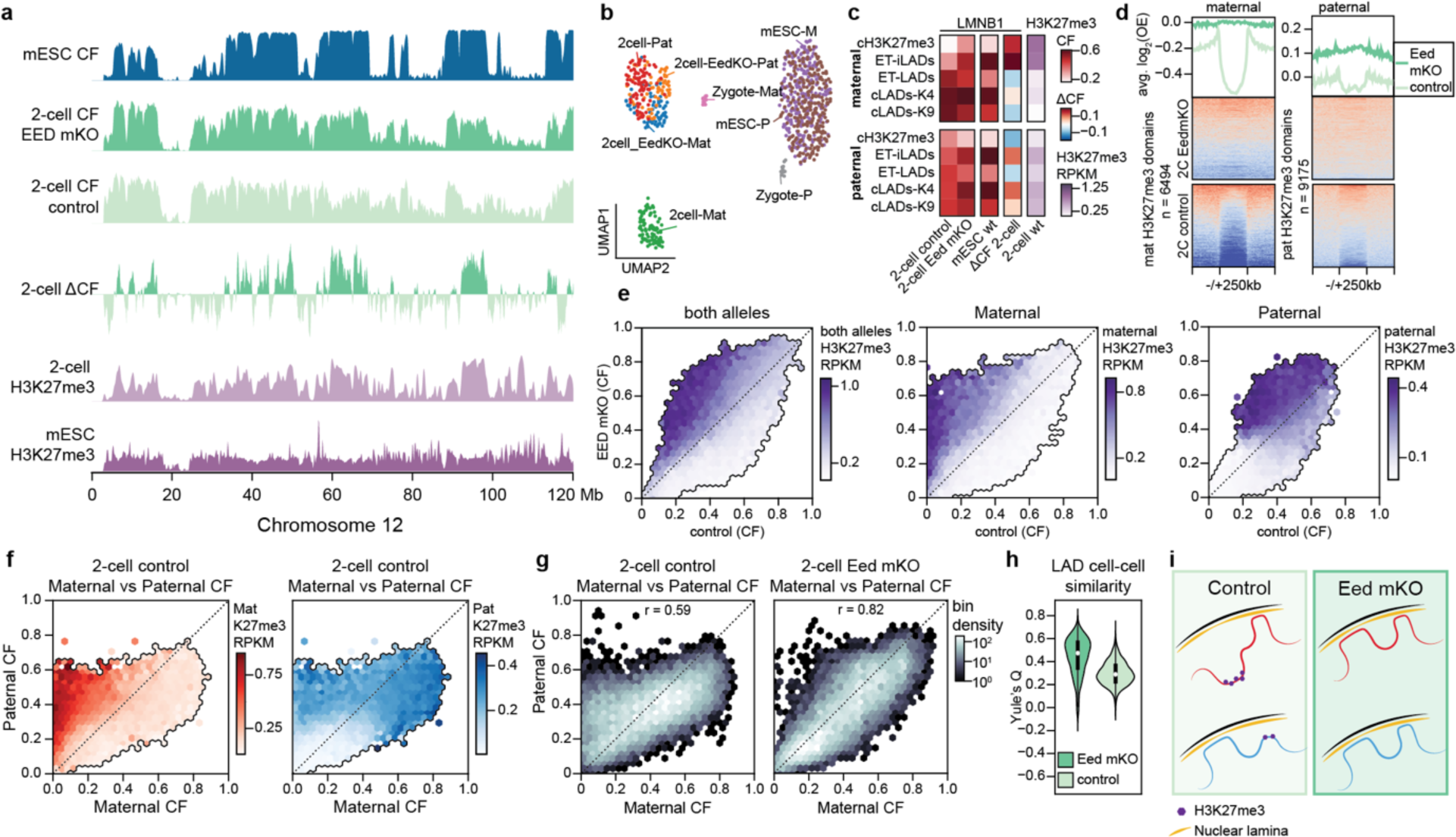
H3K27me3 antagonizes genome-NL association during early development. **a,** LAD profiles of control and *Eed* mKO 2-cell embryos, differential LMNB1 enrichment between the two conditions, mESC LAD profile and H3K27me3 profiles over chromosome 12. **b,** UMAP from all allelic-separated single-cell LAD data. Each cell is represented twice in the UMAP, once for each allele. **c,** Heatmap showing *Eed* mKO and control LMNB1 average enrichment at the 2-cell stage as well as mESC average LMNB1 enrichment and differential LAD values between the two conditions for each of the 5 genomic clusters depicted in Figure 4a. H3K27me3 average enrichment at the 2-cell stage is also plotted. **d,** Enrichment plot showing maternal (left) and paternal (right) LMNB1 enrichment at the 2-cell stage for either the *Eed* mKO or control condition over maternal H3K27me3 domains and surrounding 250 kb. Heatmaps show LMNB1 signal per domain while line plots show average enrichment per condition. **e,** Correspondence between LMNB1 values in the *Eed* mKO condition and the control condition for both alleles (left), or maternal and paternal alleles separately (right). The color scale refers to the corresponding combined or allele-specific H3K27me3 values. **f,** Correspondence between maternal and paternal Dam-LMNB1 CF. The color within the plot refers to allelic specific H3K27me3 RPKM values: maternal on the left (red) and paternal on the right (blue), **g,** Correspondence between Paternal and Maternal LMNB1 CF in control (left) or Eed mKO conditions (right) **h,** Violin plots with distribution of Yule’s Q values as a measure of LAD cell-cell similarity for *Eed* mKO (dark green) and control (light green). **i,** Model of the relationship between H3K27me3 and genome-nuclear lamina association during early development. Non-canonical broad H3K27me3 domains are present in regions that particularly in early developmental stages are not located at the nuclear lamina. Upon H3K27me3 depletion using the *Eed* mKO these regions relocate to the nuclear periphery. In addition, H3K27me3 is enriched in regions that show differential lamina association between the paternal and maternal allele. In the absence of H3K27me3 allelic LAD differences are reduced.

We then asked how the different genomic clusters identified above were affected by Eed depletion. We also included cluster 5 in this analysis. This cluster is typified by constitutive H3K27me3 enrichment throughout all stages and we therefore refer to these regions as cH3K27me3 (Figure 4a and Extended Data Figure 5b and c). Strikingly, the clusters that gained association to the nuclear lamina were, for both the maternal and paternal alleles, regions rich in H3K27me3 in the *wt* 2-cell embryo (Figure 5c). Those changes resulted in NL association values that resembled those in mESCs (Figure 5c), suggesting that the H3K27me3 absence reverts genome-NL positioning back to its canonical state.

We then called H3K27me3 2-cell maternal and paternal domains over which we plotted the corresponding *Eed* mKO and control LMNB1 enrichment values. While the control showed a clear depletion of NL association at H3K27me3 domains, the *Eed* mKO condition showed LMNB1 enrichment at these regions, indicating that for both alleles, H3K27me3 regions are relocated to the lamina in the mKO condition (Figure 5d). Similarly, genomic bins that gain NL association in the *Eed* mKO condition corresponded to H3K27me3-rich regions in the *wt* context, both for the allele-separated and non-separated data (Figure 5e). The same positive association with NL association gain was found for 2-cell H3K9me3 and mESC Dam-LMNB1 CF but not 2-cell H2AK119ub1 a histone mark deposited by Polycomb repressive complex 1 (PRC1) (Extended Data Figure 6b). This is not surprising as H2AK119ub1 has been described to progressively re-establish canonical distributions starting from the 2-cell stage, unlike non-canonical H3K27me3 which persists until implantation has occurred^11^. Together these results suggest that H3K27me3 or PRC2 play a key role in determining the atypical LAD organizations during early developmental stages.

### Allelic LAD differences are reduced in the absence of H3K27me3

Following our observation that paternal and maternal LAD differences at the 2-cell stage appeared to be reduced in the *Eed* mKO condition, we hypothesized that H3K27me3 could be related to the pronounced LAD asymmetry typical of early developmental stages^1^. Comparison of control Dam-LMNB1 values in maternal vs paternal allele showed that regions that had stronger lamina association in the paternal allele, were H3K27me3-rich in the maternal allele (Figure 5f). A similar trend was also seen in the converse situation although to a lesser extent, likely due to overall low levels of H3K27me3 (Figure 5f).

We then wondered if removing H3K27me3 would have an effect on LAD allelic differences. Visual inspection of LAD tracks indicated a reduction of LAD allelic asymmetry upon *Eed* and H3K27me3 loss (Extended Data Figure 6a and c). This was confirmed by the clear increase in correlation between allelic Dam-LMNB1 values upon *Eed* mKO (Figure 5g and Extended Data Figure 6d). This increase in correlation was only scored in regions with H3K27me3 as regions with no H3K27me3 showed no change in correlation between maternal and paternal LAD values (Extended Data Figure 6e).

Finally, we used the single-cell LAD data to determine whether H3K27me3 depletion also played a role in LAD heterogeneity across single cells of the 2-cell stage (Figure 1d). Indeed, we found that *Eed* mKO profiles showed higher Yule’s Q values compared to the control and thus higher cell-cell similarity (Figure 5h). This reduction in LAD heterogeneity between single cells upon H3K27me3 depletion was seemingly exclusive to the paternal allele. As a consequence, paternal LADs showed similar levels of cell-cell variability to the ones observed in the maternal genome (Extended Data Figure 6f).

Collectively, our results show that the non-canonical distributions of H3K27me3 prevents conventional NL-contacts from forming, contributes to paternal cell-cell LAD variability and dictates allelic LAD asymmetry during early mouse development (Figure 5i).

## Discussion

Here, we have profiled LADs across preimplantation stages and mESCs in single cells and identified a potential mechanism for the atypical distribution of the genome at the nuclear periphery during early development.

### High cell-cell variability of genome-NL contacts at the 2-cell stage

We show that the 2-cell stage genome varies extensively in its localization at the periphery of the nucleus among single cells. While some level of LAD single-cell variability is expected^18, 19^, the unusually high LAD heterogeneity in early development could be due to the totipotent nature of these stages. Interestingly, paternal zygotic LADs are not as variable, potentially due to either *de novo* establishment of LAD patterns in the absence of paternally inherited chromatin modifications or to maintenance of LAD patterns carried over from sperm, which are yet to be profiled.

Genomic regions on the centromeric end of the chromosome associated to the nuclear lamina in a high proportion of cells at the 2-cell stage, which could be a result of centromeres locating at the periphery of nucleus. 3D-FISH experiments at the late 2-cell stage report on pericentromeric repeats localizing to nucleolar precursor bodies and, although not reported, the published images also suggest proximity to the nuclear periphery^25^. Specific nuclear lamina association along chromosomes has also been described during the G1 cell cycle phase and in the context of oncogene-induced senescence although in these cases the enrichment was seen at the telomeres^26, 27^.

Genomic association to the nuclear lamina has been shown to precede chromatin topology in early development^1^. In addition, LADs are known to vary in a coordinated fashion according to DNA-DNA interactions in nuclear 3D space^19^. At the zygote stage, in a context where 3D-genome topology is mostly absent, we have shown that LADs, although present, show little coordination in their variability. A recent publication revealed single-cell structures in the paternal zygotic genome that vary extensively from cell to cell, thereby obscuring the existence of structure in the allele-combined data^28^. These single-cell domains were shown to relate to lamina association indicating that, at the zygote stage, LAD cell-cell variability and heterogeneity in zygote chromatin topology could be linked.

Variability in NL-association patterns at the 2-cell stage seem to be highest in the paternal genome. This may be due to the reduced level of histone modifications reported for the paternal allele^2^ which could reduce its affinity for specific nuclear compartments.

### Uncoupling between 3D-nuclear genome organization and gene expression at the 2-cell stage

We did not observe variability of histone PTMs and open chromatin at the same scale as variability in nuclear lamina association. However, we do not exclude that there might be heterogeneity across single cells at a smaller genomic scale (for example at the level of promoters or genes). LAD cell-cell variability having little impact on chromatin state and gene expression levels at the 2-cell stage is a very surprising finding. We hypothesize that the totipotent cells that populate the early embryo show unusual uncoupling between 3D-genome organization within the nucleus and gene regulation. It remains unclear whether regions that don’t contact the lamina locate to nuclear precursor bodies similar to what has been seen in a human cell line after cell division^29^.

### Non-LADs at the 2-cell stage are instead enriched in H3K27me3

Non-canonical H3K27me3 broad domains have been described to locate in distal regions – not in the vicinity of gene promoters - and these domains form during oogenesis persisting until post-implantation stages^14^. Here, we find that these non-canonical H3K27me3 regions have a strong correspondence with mESC LADs but show decreased NL-association at the 2-cell stage compared to mESCs or zygotic paternal genomes. Depletion of H3K27me3 via maternal KO of a component of PRC2 prompts these same regions to become LADs. Our study thus indicates that non-canonical H3K27me3 promotes dissociation from the NL during preimplantation development (Figure 5i).

Following fertilization, histone PTMs in the paternal genome are mostly established *de novo* while the maternal chromatin tends to retain a large part of the epigenetic information from the oocyte^8, 12–14^. This leads to extensive allelic epigenetic asymmetry in early developmental stages, which in the case of H3K27me3 is retained up until implantation when canonical distributions are recovered^14^. In this study, we have shown that removal of H3K27me3 in early stages reduces allelic asymmetry in genome-NL interactions at the 2-cell stage, implicating this histone PTM in establishing allelic LAD asymmetry (Figure 5i). These results demonstrate that allele-specific epigenetic features can determine distinct nuclear localizations of the maternal and paternal genomes.

H3K27me3 depletion also had an effect on cell-cell variability in genome-lamina association that is particularly high at the 2-cell stage. Strikingly, only paternal LADs seemed to become less variable. The uniformity in H3K27me3 signal observed across single cells (Figure 3a and d) is mostly a readout from the maternal allele and obscures possible H3K27me3 variability in the paternal allele. We thus speculate that the constant H3K27me3 signal on the maternal genome consistently segregates the genome into H3K27me3 domains and LADs across single cells. The low and possibly variable H3K27me3 signal on the paternal genome, on the other hand, may cause a “tug-of-war” between the antagonistic effect of H3K27me3 on lamina association and the mechanisms that drive conventional LAD formation such as A/T content and multivalency. Removal of H3K27me3 on both alleles in the *Eed* mKO embryos would thus result in the predominance of only LAD-driven mechanisms leading to more typical LAD conformations and reduced cell-to-cell heterogeneity.

## Conclusion

The study of epigenetics, nuclear organization and transcription during the first days of embryonic development has greatly benefited from the development of novel low-input technologies. These have brought much needed insight into the nuclear events that occur from the moment gametes fuse to form a totipotent zygote until implantation^9–14^. An overall view of non-canonical epigenetic features and major restructuring of genomic organization has emerged but little is known about how these processes are connected and what their role in embryonic development may be. Here, we propose a model whereby H3K27me3 antagonizes genome-NL association during preimplantation development causing both atypical NL association and allelic LAD asymmetry (Figure 5i). In support of our findings, a recent study reported increased nuclear lamina interactions of B compartment regions rich in H3K27me3 upon inhibition of another PRC2 component – Ezh2 – in K562 cells^30^. In the present study, we demonstrate that this interplay between Polycomb and genome-NL association is likely mechanistically involved in the processes that so dramatically reorganize the nuclear architecture during preimplantation development.

## Methods

### RNA synthesis

All constructs were cloned into an *in vitro* transcription vector previously described in ^1^, linearized, purified using the QIAquick PCR Purification Kit (Qiagen) and transcribed using the T3 mMessage mMachine kit (Invitrogen, AM1348) according to manufacturer instructions. The synthesized RNA was purified using the MEGAclear kit (Invitrogen, AM1908) and eluted in10 mM Tris-HCl pH 7.5 and 0.1 mM EDTA.

### Animal care and zygote injection

All animal experiments were approved by the animal ethics committee of the Royal Netherlands Academy of Arts and Sciences (KNAW) under project license AVD801002016728 and study dossiers HI173301 and HI213301. Embryos were collected from B6CBAF1/J females crossed with CAST/EiJ males for the hybrid experiments and B6CBAF1/J males for non-hybrid experiments. The *Eed* floxed mouse (*Eed^fl/fl^*) was provided by Prof. Stuart H. Orkin. To obtain *Eed* maternal KO embryos, we crossed *Eed^fl/fl^;Gdf9-icre* females (on a C57BL/6J background) with JF1/MsJ males.

For all crosses, 7- to 10-week-old females were superovulated by injecting pregnant mare serum gonadotropin (PMSG, 5IU, MSD, Cat#A207A01) and human chorionic gonadotropin (hCG, 5IU, MSD, Cat#A201A01). For *in vitro* fertilization (IVF), spermatozoa from JF1/MsJ males were capacitated in Human Tubal Fluid medium (Merck Millipore, Cat#MR-070-D) supplemented with 10 mg ml^−1^ Albumin (Sigma, Cat#A-3311) (HTF-BSA) for 1h preceding insemination. MII oocytes were collected in the insemination medium (HTF-BSA) and capacitated spermatozoa were added for fertilization. The insemination starting time point was termed as 0 hours post-fertilization (hpf).

When using standard mating, zygotes were injected about 24h post-hCG.

For both IVF and normal mating mRNA was microinjected in the cytosol of the zygote at 10hpf. A full description of constructs used, corresponding concentrations and induction conditions is described in Extended Data Table 1.

Injected zygotes were cultured in KSOM for hybrid crosses (Sigma, Cat#MR-106-D) or M16 medium (Sigma, Cat#M7292) for non-hybrid crosses covered with mineral oil (Sigma Cat# M8410) at 37°C with 5% CO_2_ and 5% O_2_ air.

To increase the quality of DamID signal, untethered Dam and Dam-Cbx1 constructs were fused to an ERT2 domain so that the fusion protein would be translocated to the nucleus upon 4-Hydroxytamoxifen addition (4-OHT, Sigma, Cat#SML1666) (Extended Data Table 1).

### Embryo collection and dissociation and scDam&T-seq processing

Embryos were collected by mouth pipetting at 29-31 hours post-hCG for the zygote stage, 52-55 hours post-hCG for the 2-cell stage and 75-78 hours post-hCG for the 8-cell stage (Extended Data Table 1). The zona pellucida was removed using Tyrode’s acid (Sigma, Cat#T1788), washed in M2 medium (Sigma, Cat#MR-015) and placed in TryplE (Gibco, Cat#12605010) where embryos were dissociated into single cells one by one and placed in M2 medium before single-cell collection into a 384-well plate containing 5uL of mineral oil and 100nL of barcoded polyadenylated primers. All scDam&T-seq steps were performed as previously described^31^. Briefly, cells were lysed and reverse transcription was performed followed by second-strand synthesis in order to convert the RNA of the cell into cDNA. After a proteinase K step, methylated GATCs resulting from Dam enzyme activity were specifically digested with DpnI and double-stranded adapters containing cell-specific barcodes were ligated. At this point, cells with non-overlapping barcodes were pooled together to undergo in vitro transcription which amplifies both the transcriptional and genomic product in a linear manner due to a T7 promoter to both the double-stranded DamID adapters and the polyadenylated primers. The resulting amplified RNA (aRNA) was reverse transcribed and library preparation was performed as previously described^32^. Libraries were sequenced on the Illumina NextSeq500 (75-bp reads) or NextSeq2000 (100-bp reads) platform. For scDam&T-seq processing of mESC cells specifically, half volumes were used in all reactions to reduce overall processing costs.

### Cell culture

Cell lines were grown in a humidified chamber at 37 °C in 5% CO_2_ and were routinely tested for mycoplasma. Mouse F1 hybrid Cast/EiJ (paternal) x 129SvJae (maternal) embryonic stem cells (mESCs; a gift from the Joost Gribnau laboratory) were cultured on 6-well plates with irradiated primary mouse embryonic fibroblasts (MEFs) in mESC culture media (CM) defined as follows: Glasgow’s MEM (G-MEM, Gibco, 11710035) supplemented with 10% FBS, 1% Pen/Strep, 1x GlutaMAX (Gibco, 35050061), 1x MEM non-essential amino acids (Gibco, 11140050), 1 mM sodium pyruvate (Gibco, 11360070), 0.1 mM β-mercaptoethanol (Sigma, M3148) and 1000 U/mL ESGROmLIF (EMD Millipore, ESG1107). mESCs were alternatively cultured in feeder-free conditions on gelatin coated plates (0.1% gelatin, in house) in 60%-BRL medium, defined as a mix of 40% CM medium (as defined) and 60% conditioned CM medium (incubated 1 week on Buffalo Rat Liver cells), supplemented with 10% FBS, 1% Pen/Strep, 1x GlutaMAX, 1x MEM non-essential amino acids, 0.1 mM β-mercaptoethanol and 1000 U/mL ESGROmLIF. Cells were split every 2-3 days and medium was changed every 1-2 days. This mESC cell line does not contain a Y chromosome.

### Generation of mouse embryonic stem cell lines

The stable clonal F1 hybrid mESC line expressing the Dam-LaminB1 fusion protein was generated from an EF1α-Tir1-IRES-neo expressing mother line (generated with lentiviral transduction)^18^. The Dam construct was CRISPR targeted into this line by knocking in mAID-Dam in the N terminus of the LMNB1 locus. The donor vector (designed in house, generated by GeneWiz) carried the Blasticidin-p2A-HA-mAID-Dam cassette, flanked on each side by 1000-bp homology arms of the endogenous LMNB1 locus (pUC57-BSD-p2A-HA-mAID-Dam). The Cas9/guide vector was the p225A-LmnB1-spCas9-gRNA vector, with a guide RNA inserted to target the 5’UTR of the LMNB1 locus (sgRNA: 5’ CACGGGGGTCGCGGTCGCCA 3’). For transfections in general, cells were cultured on gelatin-coated 6-well plates in 60% BRL-medium at 70%–90% confluency. Cells were transfected with Lipofectamin2000 (Invitrogen, 11668030) according to the supplier protocol with 1.5 µg donor vector and 1.5 µg Cas9/guide vector. At 24 hours after transfection, GFP positive cells were sorted on a BD FACsJazz Cell sorter and seeded on gelatin-coated plates in 60% BRL-medium. 48 hours after sorting, cells were started on antibiotic selection with 60% BRL-medium containing 3.0 µg/mL Blasticidin (ThermoFisher, A1113903) and 0.5 mM indole-3-acetic acid (IAA, Sigma, I5148) and cells were refreshed every 2-3 days. From this point onwards, 0.5 mM IAA is added to the medium during normal culturing conditions to degrade the mAID-Dam-Lamin B1 fusion protein via the auxin protein degradation system^33^. After 6 days of antibiotic selection, single cells were sorted into 96-well plates containing MEFs using the BD FACsInflux Cell sorter and grown without antibiotic selection in CM medium with 0.5 mM IAA. Clones grew out in approximately 10 days and were screened for correct integration by PCR with primers from Dam to the LMNB1 locus downstream of targeting construct; fw-TTCAACAAAAGCCAGGATCC and rev-TAAGGAATCTGGTGCACAGAACACC. The heterozygous expression of the Dam-Lamin B1 fusion protein was further confirmed by Western blot using an anti-HA antibody at 1 in 5000 dilution (Abcam, ab9110) and an anti-LaminB1 antibody at 1 in 5000 dilution (Abcam, ab16048). To prevent silencing of the EF1α-Tir1-IRES-neo construct due to the flanking lentiviral construct sequences, the Tir1 construct was additionally knocked-in into the TIGRE locus using CRISPR targeting. This integration was generated by co-transfection of the donor vector pEN396-pCAGGS-Tir1-V5-2A-PuroR TIGRE (Addgene plasmid, #92142) and Cas9-gRNA plasmid pX459-EN1201 (backbone from Addgene plamid #62988, guide from Addgene plasmid #92144^34^, sgRNA: 5’ ACTGCCATAACACCTAACTT 3’). Cells were transfected with Lipofectamine3000 (ThermoFisher, L3000008) according to the supplier protocol with 2 µg donor vector and 1 µg Cas9-gRNA vector. At 24 hours after transfection, GFP positive cells were sorted on a BD FACsJazz Cell sorter and seeded on gelatin-coated plates in 60% BRL-medium. 48 hours after sorting, cells were started on antibiotic selection with 60% BRL-medium containing 0.8 µg/mL Puromycin (Sigma, P9620) and 0.5 mM IAA. Cells were refreshed every 2-3 days and selected for 5-10 days. The Tir1-puro clones were screened for the presence of Tir1 by PCR from the CAGG promoter to Tir1 with the primers fw-CCTCTGCTAACCATGTTCATG and rev-TCCTTCACAGCTGATCAGCACC, followed by screening for correct integration in the TIGRE locus by PCR from the polyA to the TIGRE locus with primers fw-GGGAAGAGAATAGCAGGCATGCT and rev-ACCAGCCACTTCAAAGTGGTACC. The Tir1 expression was further confirmed by Western blot using a V5 antibody (Invitrogen R960-25). Upon further characterization of the best clone, a 70-bp deletion was found directly after the transcription start site the wildtype LMNB1 allele, causing frameshift, which was most likely the result of the CRISPR targeting. Cell viability and growth rates were not visibly affected. This deletion was repaired using 200 bp ssDNA utramere oligo’s (IDT) with 65 bp homology arms on each side of the deletion as donor and a p225A-LmnB1-repair-spCas9-gRNA vector (sgRNA: 5’ GCGGGGGCGCTACAAACCAC 3’). Cells were transfected with Lipofectamine 3000 according to the supplier protocol with 1.5 µg donor oligo and 1 µg Cas9-gRNA vector. At 24 hours after transfection, GFP positive cells were sorted into 96-well plates containing MEFs using the BD FACsJazz Cell sorter and grown without antibiotic selection in CM medium. Clones were screened for correct repair of the wildtype LMNB1 allele by PCR around the original deletion with the primers fw-ACTCACAAAGGGCGTCTGGC and rev-GTGACAATCGAGCCGGTACTC. Correct expression of the mAID-Dam-Lamin B1 fusion protein as well as the wildtype Lamin B1 protein was confirmed using Immunofluorescence staining using an anti-HA antibody at 1 in 500 dilution (Cell Signaling Technologies, C29F4) and an anti-LaminB1 antibody at 1 in 500 dilution (Abcam, ab16048), followed by confocal imaging. All successfully repaired clones were subsequently screened for their level of induction upon IAA removal by m6A-PCR, evaluated by gel electrophoresis^6, 35^, followed by DamID2 sequencing in bulk^31, 35^, to select a heterozygous clone with a correct karyotype with the best signal-to-noise ratio of enrichment over LAD regions. This clone is labelled as F1ES mAID-Dam-LaminB1 #2B4.

### mESC Dam-Lamin B1 induction and FACS sorting for single-cell experiments

Expression of the mAID-Dam-Lamin B1 fusion protein in the F1ES cell line was suppressed by addition of 0.5 mM IAA during standard culturing. When plated for scDam&T-seq experiments, the cells were passaged at least two times in feeder-free conditions on 6-well plates coated with 0.1% gelatin in 60%-BRL medium. Cells were kept at 1 mM IAA for the final 48 hours before the start of the experiment. 6 hours before harvesting of cells, the IAA was removed by washing three times with PBS and refreshing with 60%-BRL medium without IAA. FACS was performed on BD FACSJazz or BD FACSInflux Cell Sorter systems with BD Sortware. mESCs were harvested by trypsinization, centrifuged at 300 g, resuspended in 60%-BRL medium containing 10 mg/mL Hoechst 34580 (Sigma, 63493) per 1×10^6^ cells and incubated for 45 min at °C in 5% CO_2_. Prior to sorting, cells were passed through a 40-mm cell strainer. Propidium iodide (1 mg/mL) was used as a live/dead discriminant. Single cells were gated on forward and side scatters and Hoechst cell cycle profiles. Index information was recorded for all sorts. One cell per well was sorted into 384-well hard-shell plates (Biorad, HSP3801) containing 5 µL of filtered mineral oil (Sigma, 69794) and 50 nL of 1.5 mM barcoded CEL-Seq2 primer^18, 31^.

### Processing of scDamID and scDam&T-seq data

Data generated by the DamID and scDam&T-seq protocols was largely processed with the workflow and scripts described in Markodimitraki et al. (2020)^31^(see also www.github.com/KindLab/scDamAndTools). The procedure is described in short below.

#### Demultiplexing

All reads are demultiplexed based on the barcode present at the start of R1 using a reference list of barcodes. In the case of scDam&T-seq data, the reference barcodes contain both DamID-specific and CEL-Seq2-specific barcodes. In the case of the scDamID data, the reference barcodes only contain DamID-specific barcodes. Zero mismatches are allowed between the observed barcode and reference. The UMI information, also present at the start of R1, is appended to the read name.

#### DamID data processing

DamID reads are aligned using bowtie2 (v. 2.3.3.1)^36^ with the following parameters: “--seed 42 -- very-sensitive -N 1” to the mm10 reference genome. In the case of paired-end data (scDam&T-seq), only R1 is used as that contains the digested GATC site. The resulting alignments are then converted to UMI-unique GATC counts by matching each alignment to known strand-specific GATC positions in the reference genome. Any reads that do not align to a known GATC position or have a mapping quality smaller than 10 are removed. Up to 4 unique UMIs are allowed for single-cell samples to account for the maximum number of alleles in G2. Finally, counts are binned at the desired resolution.

#### CEL-Seq2 data processing

CEL-Seq2 reads are aligned using hisat2 (v. 2.1.0)^37^ with the following parameters: “--mp ‘2,0’ --sp ‘4,0’”. For the alignment, only R2 is used, as R1 contains the sample barcode, UMI and poly-A tail, which have already been processed during demultiplexing. As reference, the mm10 reference genome and the GRCm38 (v. 89) transcript models are used. Alignments are subsequently converted to transcript counts per gene with custom scripts that assign reads to genes similar to HTSeq’s^38^ htseq-count with mode “intersection_strict”.

#### Allele-specific alignment of DamID and CEL-seq2 reads

In the case of samples derived from hybrid crosses, we used strain-specific SNPs to assign reads to a parent. For this, we obtained SNP information from the Mouse Genomes Project of the Sanger Wellcome Institute for all used strains except for JF1/Ms, which were obtained from the MoG+ website of the RIKEN BioResource Center (https://molossinus.brc.riken.jp/mogplus/#JF1). These SNPs were subsequently substituted in the mm10 reference genome to generate strain-specific reference files. DamID and CEL-seq2 reads were subsequently aligned to the reference files of both strains as described above. Since all hybrid data was generated with scDam&T-seq, paired-end data was available for the DamID readout and both R1 and R2 were used in aligning to maximize SNP coverage. Using a custom script, the alignments of each read to the two genotypes were subsequently evaluated w.r.t. number of mismatches and alignment score. The read was then attributed to the better scoring genotype. In the case of a tie (i.e. equal number of mismatches and same alignment score), the read was considered to be ambiguous. This procedure results in three files for each sample: one alignment file for each genotype and one file with ambiguous reads. For the samples derived from the B6CBAF1/J x CAST/EiJ cross, SNPs from three different backgrounds can be present: CBA/J and C57BL/6J SNPs from the B6CBAF1/J mother and CAST/EiJ from the father. Reads derived from this cross were thus aligned to the three reference genomes representing these strains and split based on their alignment scores as described. Reads attributed to CBA/J and/or C57BL/6J were considered as maternal reads, reads attributed to CAST/EiJ as paternal reads, and reads tying between CAST/EiJ and CBA/J or C57BL/6J as ambiguous.

#### Processing allele-specific DamID and CEL-seq2 read to UMI-unique counts

For both CEL-seq2-derived and DamID-derived reads information from R2 is used to attribute them to a genotype. However, in both cases, the IVT and fragmentation steps in the scDam&T-seq protocol can result in copies of the original mRNA/DNA molecule of different lengths and thus different R2 sequence content. As a result, different copies of the same molecule sometimes overlap SNPs and sometimes do not. For this reason, it is import to perform UMI flattening per gene or GATC position for all alignment files (both genotypes and ambiguous) simultaneously. Per gene or GATC position, only one unique UMI is allowed across the genotypes. If a UMI was observed for one genotype and in the ambiguous reads, the unique count was attributed to the genotype. If a UMI was observed for both genotypes, the unique count was considered to be ambiguous. For this procedure, modified versions of the DamID and CEL-seq2 counting scripts were used that consider all three alignment files in parallel. Counting was otherwise performed as described above.

#### Filtering of DamID data

Samples were filtered w.r.t. their DamID readout based on the number of observed unique GATCs and their information content (IC). The IC is a measure for the amount of true signal present in a sample relative to the amount of background. The background is determined based on a comparison of the observed signal with the density of mappable GATCs in the genome. The procedure is explained in detail in Rang and de Luca (2022)^22^ and the code can be found on GitHub (https://github.com/KindLab/EpiDamID2022). Since the fraction of the Dam-methylated genome varies per Dam-construct and per embryonic stage, the thresholds for the number of unique GATCs and IC were fine-tuned per dataset (Extended Data Table 2).

In the case of samples derived from hybrid crosses, DamID data was additionally filtered based on the presence of both a maternal and paternal allele. In particular, at least 25% of allele-specific counts should come from each parent. In practice, this resulted in the removal of samples that exclusively had maternal-derived material, likely due to the presence of unfertilized oocytes undergoing spontaneous parthenogenesis. We observed no samples containing >75% paternal-derived material. For analyses using data of the combined alleles the same filtering on unique GATCs and IC was applied. For analyses using allele-specific data, only samples were used that had a total number of allele-specific GATC counts equal to the general depth threshold of that condition. We performed this additional select to prevent high levels of noise due to sparsity in allele-specific data. The numbers of cells that passed the aforementioned thresholds are documented in Extended Data Figures 1c, 3c and 4b and unique number of GATC distribution of DamLMNB1-expressing cells that passed those thresholds is illustrated in Extended Data Figure 1d.

For genome-wide analyses, we additionally performed filtering on the genomic bins that were included. For analyses that were not allele-specific, we excluded all genomic bins that contain fewer than 1 mappable GATC per kb. For allele-specific analyses, we additionally removed bins for which less than 10% of the contained GATCs could be attributed to an allele. In addition, we removed bins for which we empirically observed that 98% of allele-specific DamID counts were attributed to only one allele.

#### Filtering of CEL-seq2 data

Samples were filtered w.r.t. their CEL-seq2 readout based on the observed number of unique transcripts (≥3,000), the percentage of mitochondrial transcripts (<15%), and the percentage of ERCC spike-in derived reads (<0.5%). For all stages and constructs these thresholds were the same. In addition, hybrid samples that were suspected to have undergone spontaneous parthenogenesis based on their DamID readout (see above) were also excluded from transcriptional analyses. This filtering could not be performed based on the transcriptional readout, since the vast majority of transcripts at the zygote and 2-cell stage are maternally contributed.

#### Computing DamID binary contacts

Single-cell count tables were further processed to binary contacts, which give an indication for each genomic bin weather a sample had an observed contact with the Dam construct. To determine binary contacts, samples were binned at 100,000-bp resolution and depth normalized by log 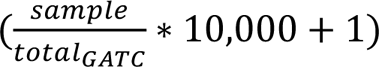. Allele-specific files were normalized for the total number of counts attributed to *either* allele. Subsequently, the samples were smoothened with a Gaussian kernel (s.d. 150 kb). Both the large bin size and smoothing minimize noise that may occur in sparse single-cell samples. The observed signal was subsequently compared with a control, which has been depth normalized and smoothed in a similar manner. In the case of the Dam-LMNB1 and Dam-only constructs, the control is the density of mappable GATCs. In the case of Dam-H3K27me3, Dam-H3K27me3mut, and Dam-Cbx1, the control is the average single-cell signal of all Dam-only samples of the same embryonic stage, since these constructs are free-floating in the nucleus and are more prone to accessibility biases^22^. Contacts were then called when the difference between the observed signal and the control was bigger than 0 for Dam-LMNB1 and Dam-only, or bigger than log(1.1) ≈ 0.095 for the remaining constructs.

#### Contact frequency and in silico population profiles

Contact frequency (CF) was determined for each genomic bin as the fraction of single-cell samples with an observed contact in that bin. As a result, CF values range between 0 and 1. Since binary contacts are only determined at a 100-kb resolution, CF profiles are only available at this resolution. For analyses requiring higher resolutions, we generated *in silico* population profiles by combining the count data of all single-cell samples per condition. The *in silico* population profiles were subsequently depth normalized (RPKM) and normalized for a control. Allele-specific files were depth normalized for the total number of counts attributed to *either* allele. In the case of Dam-LMNB1 and Dam-only, the control is the density of mappable GATCs. For the other constructs, the control is the *in silico* population data of the Dam-only samples.

### Processing of published data

Accession numbers of all public datasets used are described in Extended Data Table 3.

#### ATAC-seq, ChIP-seq, and CUT&RUN

Reads were aligned using bowtie2 (v. 2.3.3.1) with the following parameters: “--seed 42 -- very-sensitive -N 1”. For paired-end ATAC-seq files, the following additional parameters were used: “-X 1000 --no-mixed --no-discordant”. Indexes for the alignments were then generated using “samtools index” and genome coverage tracks were computed using the “bamCoverage” utility from DeepTools (v. 3.3.2)^39^ with the following parameters: “--ignoreDuplicates -- minMappingQuality 10”. For samples derived from hybrid crosses, a similar strategy was used as for our own scDam&T-seq data to attribute reads to alleles: SNPs were incorporated into the mm10 reference genome to generate parental-specific references. Reads were aligned to both genomes, after which reads were attributed to a specific parent or ambiguous alignment files based on the number of mismatches and alignment scores. These three alignment files were then separately processed with DeepTools.

#### DamID

Data from our previous study (Borsos et al., 2019; available on GEO under GSE112551) was reprocessed using the same procedures as used for the current data. Since this data was generated with the first version of the scDamID protocol^19^, no UMIs are present to identify PCR duplicates. To limit amplification artefacts only 1 count per strand-specific GATC position was maintained.

#### Hi-C

Published Hi-C data was obtained from GEO (GSE82185) and processed from raw sequencing reads to interaction matrices using Hi-C Pro (v. 2.11.4) using the recommended workflows for non-allelic and allele-separated data. The obtained interaction matrices were subsequently converted to “.cool” format using the “hicConvertFormat” command from HiCExplorer (v. 2.2.1.1). The matrices were subsequently normalized and corrected for biases using the “cooler balance” functionality from Cooler (v. 0.8.11)^40^. Further processing and visualization of the normalized Hi-C matrices was performed using CoolTools (v. 0.5.1)^41^. Compartment scores were computed with cooltools using normalized interaction matrices at a resultion of 100 kb.

#### Methyl-seq

Processed files were downloaded from GEO (GSE56697). To generate binned genomic tracks, the average fraction of methylated CpGs was computed.

### Single-cell DamID analyses

#### Single-cell DamID UMAP

The UMAPs based on the single-cell DamID readout in Figure 1b, 5b and Extended Data Figure 3d were generated by performing a PCA on the data and selected the top PCs based on the explained variance ratio (PC1-10). These PCs were used as an input to compute the UMAP. In the case of Figure 5b and Extended Data Figure 3d, the maternal and paternal readouts of all samples were treated as separate samples. Consequently, each cell appears twice in the UMAP: once with the maternal readout and once with the paternal readout.

#### Cell-cell similarity

Cell-cell similarity between cells was computed based on the binary contact data of all autosomal chromosomes. We used Yule’s Q as a metric of similarity between cells: 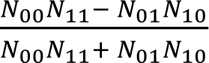, where *N*_11_ is the number of genomic bins where both samples had a contact, *N*_00_ the number of bins where neither sample had a contact, and *N*_01_ and *N*_10_ the number of bins where one sample had a contact and the other did not. Since different experimental conditions (i.e. different embryonic stages or different Dam constructs) can have different data quality (i.e. higher coverage or depth, higher signal-to-noise ratio), we devised a control for each condition that simulated the expected Yule’s Q scores based on technical variability alone. For this, we combined the data of each condition per batch (i.e. per sequencing library) and subsequently subsampled the data to generate mock single-cell samples with a number of unique GATCs equal to the actual single-cell data. We subsequently normalized and binarized the simulated single-cell data in an identical manner to the original data. The simulated dataset should thus represent samples that display the same level of technical variability (i.e. due to differences in depth and signal quality) as the original sample, without showing any true biological variation. As for the original data, we computed the Yule’s Q between all possible sample pairs to get a quantification of the cell-cell similarity. In Extended Data Figures 1e and 4e, the results for the original and simulated datasets are shown. The difference in mean between the two datasets was tested with Wilcoxon’s Rank Sum test (Extended Data Table 4). In Figures 1d and 3d, the differential Yule’s Q of all sample pairs is shown. No simulated dataset was used for the comparison between alleles (Figure 2c), since the data of the two alleles is derived from the same sample and thus has the same technical characteristics.

#### Contact run-length analysis

The contact run-length analysis was performed as in Kind et al. (2015)^19^. In short, for each cell all stretches of continuous contacts were identified based on the 100-kb binarized data. Subsequently, the number of stretches of at least length *L* were determined for *L* ranging from 1 to the maximum contact length. The average frequency of contact run-lengths was then computed per stage, as well as the standard deviation. As a control, we compared the observed contact run-lengths to those observed in the shuffled binary contact tables. This shuffling was performed in such a way that both marginals (i.e. the CF of each 100-kb bin and the total number of contacts in each cell) remained intact, using a published algorithm^42^.

#### Bin-bin coordination matrices

To quantify the coordinated association of genomic bins with the NL, we computed the Pearson correlation between all pairs of genomic bins, as previously described^19^. Since these bin-bin correlations are influenced by several factors, including the number of contacts per cell and contact frequency, we once again employed randomly shuffled binary matrices as a control (see above). For each condition, we performed 1,000 randomizations of the binary contact matrix and computed the bin-bin correlations of the resulting matrices. The observed mean and standard deviation of the correlation matrices were then used to standardize the true bin-bin correlation matrix.

#### Clustering of genomic bins

Datasets used for genomic bin clustering (100 kb) was based on the datasets described in Extended Data Table 5.

Only data of autosomal chromosomes was included. Furthermore, genomic bins were excluded from the analyses if they did not meet all of the following criteria: 1) contains at least 1 mappable GATC per kb; 2) for all allele-specific samples, at least 10% of all obtained reads/counts could be attributed to either the maternal or paternal allele; 3) for the DamID data, at least 2.5% of the observed allele-specific data of each hybrid cross can be attributed to each allele; 4) does not overlap a region annotated in the ENCODE mm10 blacklist as a “High Signal Region”; 5) does not overlap for >10% regions annotated in the ENCODE mm10 blacklist as “Low Mappability”. Criterium 1 ensures reliable DamID data; criteria 2 and 3 ensure that allele-specific resolution can be achieved; criterium 4 and 5 ensure that ChIP-seq data is reliable. For the autosomal chromosomes, this left ∼76.8% of the genomic bins to be included in the clustering.

Allele-specific LMNB1 DamID data is normalized for a control (density of allelically separable GATC motifs) that automatically corrects for biases in signal strength that are due to differential SNP density in the genome. However, allele-separated ChIP-seq data lacking an input-control are not corrected for this. As a result, allele-specific ChIP-seq RPKM values will be artificially higher in regions with high SNP density, as a higher fraction of reads could be attributed to an allele. To prevent these biases from influencing the clustering, the allele-specific RPKM values were normalized for the total fraction of reads in that bin that could be attributed to an allele. After this correction, the data of all samples were subsequently standardized and values were clipped to a range from -2.5 to 2.5.

Prior to clustering, a PCA was performed to remove redundancy in the data. The top PCs were selected based on the explained variance ratio (PC1-5), which collectively accounted for 80.5% of variance in the data. These PCs were subsequently used to compute UMAPs representing the genomic bins, as well as for K-means clustering of all bins. For the K-means clustering, a number of 8 clusters was chosen. Decreasing the number of clusters resulted in the merging of distinct clusters, while increasing the number of clusters resulted in two or more clusters with very similar behaviors.

#### Visualization of clustering

Visualization of data in the different clusters as seen in Figure 4a and Extended Data Figure 5b and c was performed as follows. For the full bin clustering (Extended Data Figure 5b and c), the datasets were grouped into the 8 different clusters. Allele-specific signal without a control dataset was corrected for biases caused by SNP density as described above. For some samples, the allelic separation of signal was very poor in a subset of bins, resulting in invalid or unreliable values in these bins. Therefore, we replaced the values of bins in which <10% of reads could be attributed to an allele with the average signal of that cluster. This mostly affected the ATAC-seq data, as most other samples were either included in the clustering (and thus bins with poor allelic separation were filtered out) and/or were based on crosses with more homogenous SNP coverage across the genome. For the heatmaps showing the average signal per cluster (Figure 4a), the average of all genomic bins (with sufficient allele separation) was taken for each cluster.

### Single-cell transcription analyses

#### Single-cell CEL-seq2 UMAP

To generate the transcriptional UMAP (Figure 1c), single-cell transcript tables were processed in R (v. 4.1.2) using Seurat (v. 4.1.0)^43^. Only samples passing transcription thresholds were included; only scDam&T-seq LMNB1 samples of embryos from homozygous crosses and the mESC samples were used. Genes with counts observed in fewer than 10 cells were excluded and data was normalized using the “NormalizeData” and “ScaleData” commands. The UMAP was then generated using the “FindVariableFeatures”, “RunPCA” and “RunUMAP” commands.

#### EED mKO vs WT differential expression

The transcriptional data of *EED* mKO and WT 2-cell embryos was processed separately for the maternally-assigned transcript counts, and the paternally-assigned transcript counts. In both cases, the data was processed using Seurat’s “NormalizeData” and “ScaleData” commands and the total number of allele-assigned reads was used to normalize the data of each allele, rather than the total maternal or paternal reads. Batch correction was performed using the “RunPCA”, “FindIntegrationAnchors”, and “IntegrateData” commands, where each individual sequencing run was considered a batch. The integrated data was then once again scaled and a PCA was performed. UMAPs were generated using the “RunUMAP” command. Differentially expressed genes between the *EED* mKO and control embryos were identified using the “FindMarkers” command. Only genes with an adjusted p-value < 0.05 were considered as differentially expressed.

#### Transcription of genes in cells with versus without NL association

To determine the effect of NL association on the expression of a gene, we determined for each gene the group of cells in which the 100kb genomic bin containing the gene TSS was in contact with the NL (“contact”) and the group of cells in which it was not (“free”). This was done separately per embryonic stage. The transcript counts of the gene and the total transcript counts were then combined for the two groups, and the expression value (as log (*RPM* + 10) was determined for each group. Genes were excluded from the analysis if either the contact or free group contained fewer than 10 cells; if the gene was expressed in fewer than 10 cells across both groups; if the gene was located on chrX or chrY; or if the gene was annotated as a maternal mRNA transcript by Park et al. (2013)^44^. In the case of allele-specific data, genes were also excluded if their TSS fell within a genomic bin that did not show sufficient allele separation (see above, *Filtering of DamID data*). The correlation in gene expression values between contact and free states was computed using Spearman’s correlation.

## Extended Data Figures

**Extended Data Figure 1.**
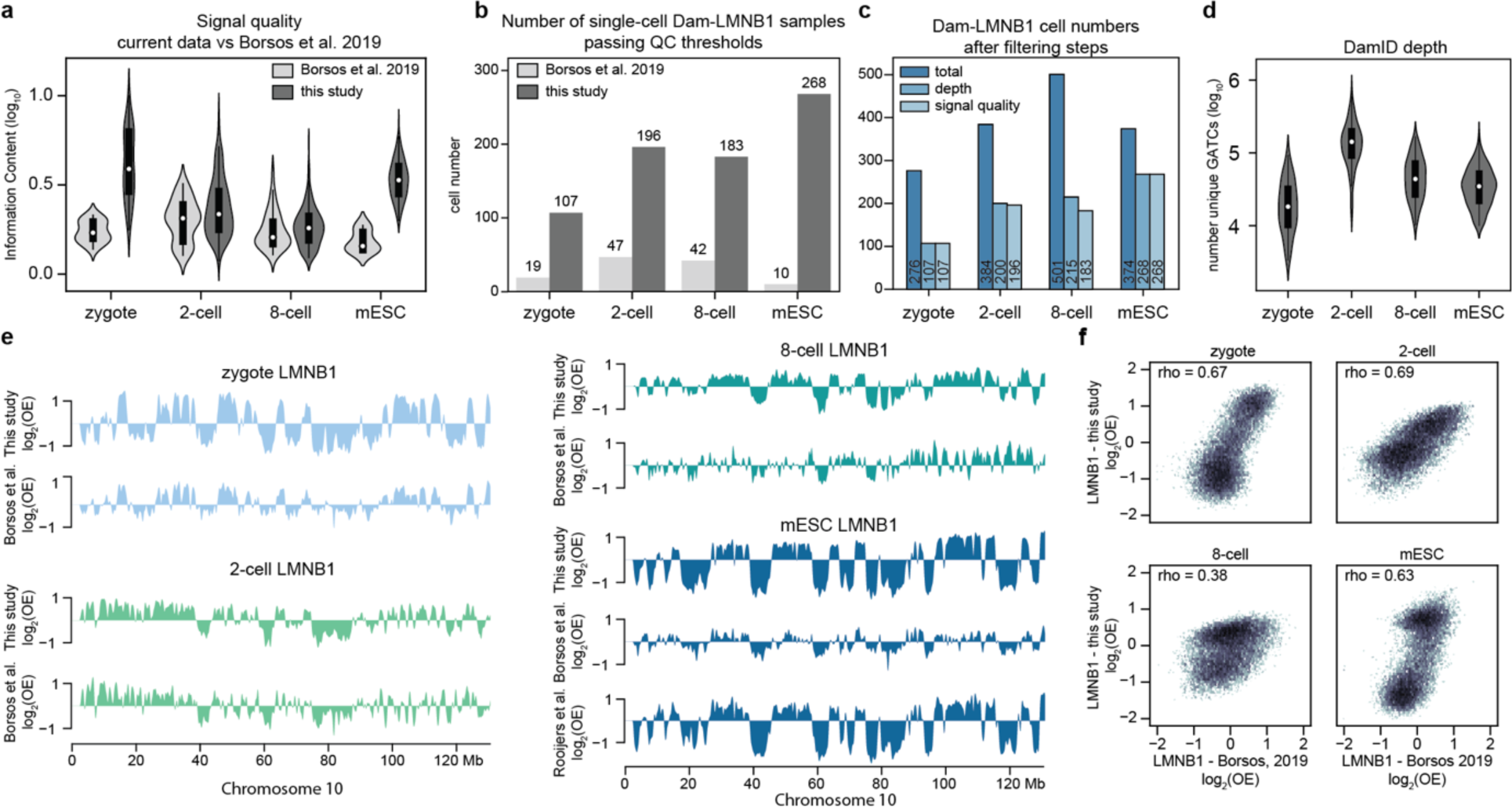
Validation of single-cell LAD data during preimplantation stages. **a,** Information content, a measure of signal quality plotted for both for the present study and Borsos et al. (2019)^1^. **b,** Comparison of cell number that pass quality thresholds for the present study and Borsos et al. (2019)^1^. **c,** Cell numbers that pass sequential filtering steps: unique number of GATCs and signal quality. **d,** Violin plot depicting the distribution of the number of unique GATCs per stage. **e,** Single-cell average LAD profiles normalized to mappability (log_2_OE) of this work compared to previous studies over the entire chromosome 10. **f,** Comparison between single-cell averages from our study and Borsos et al, 2019 for each stage using mappability normalized values (log_2_(OE)). Correlation value was obtained by calculating the Spearman rho.

**Extended Data Figure 2.**
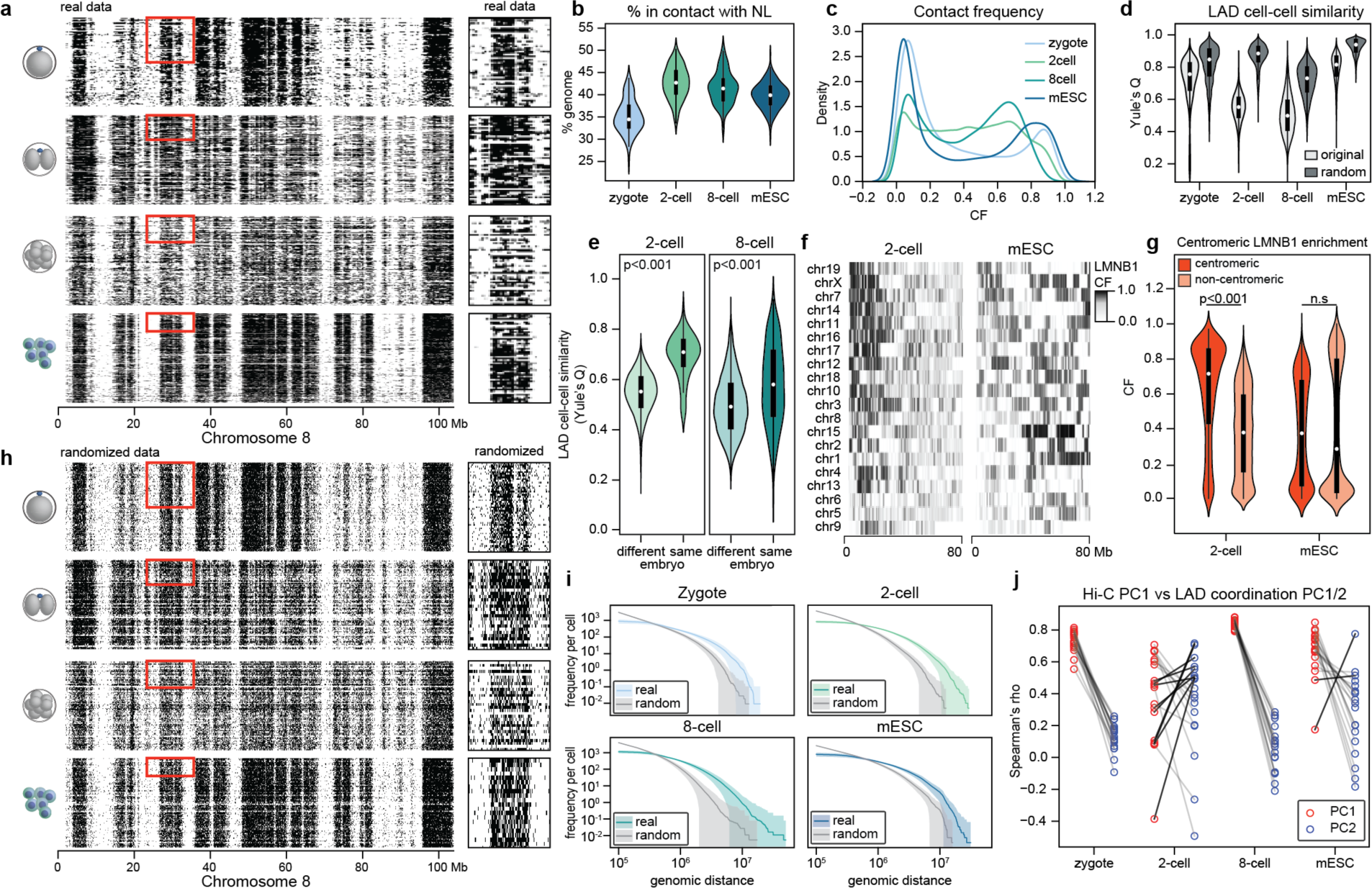
Analysis of cell-cell LAD variability in preimplantation stages and mESCs. **a,** Heatmap with single-cell binarized profiles of all cells that passed quality thresholds along the entire chromosome 8 (left panel) ordered by unique number of GATCs (DamID depth) and grouped by stage. On the right side is a magnification of the region highlighted with a red rectangle in the left heatmap which captures the same number of cells per stage. **b,** Percentage of the genome in contact with the nuclear lamina (NL) at each stage. **c,** Distribution of contact frequency (CF) per stage. **d,** Violin plot showing Yule’s Q for real data (depicted in a) and randomized data (depicted in h) per stage. Two-sided Wilcoxon rank-sum test was performed (zygote, n=107; 2-cell, n=196; 8-cell, n=183; mESCs=268) (Extended Data Table 4). **e,** Violin plot showing cell-cell similarity using Yule’s Q on cell pairs originating from the same embryo or from a different embryo at the two-cell stage (left) and at the eight-cell stage (right). Two-sided Wilcoxon rank-sum test was performed (2-cell different embryo, n=19025; 2-cell same embryo, n=85, p-value=3.36×10^-35^; 8-cell different embryo, n=14377, 8-cell same embryo: n=2276, p-value=8.00×10^-30^). **f,** Heatmap with CF distribution along the first 80 Megabases (Mb) of each chromosome for both 2-cell stage (left) and mESCs (right). Chromosomes are ordered by CF value of the first 10 Mb at the 2-cell stage. **g,** Violin plot depicting CF values in the first 30 Mb versus the remaining of the chromosome for the 2-cell stage and mESCs. Two-sided Wilcoxon rank-sum test was performed (centromeric bins, n=570, p-value<2.225×10^-308^; non-centromeric bins, n=24048, p-value=0.0013). **h,** Heatmap with single-cell binarized profiles of randomized data based on stage CF and unique number of GATCs per cell in (a) along the entire chromosome 8 (left panel) ordered by unique number of GATCs (DamID depth) and grouped by stage. This randomized data allows to control for variability originating from differences in signal quality across stages. On the right side is a magnification of the region highlighted with a red rectangle in the left heatmap which captures the same number of cells per stage. **i,** Distribution of Dam-LMNB1 contact run lengths in single-cell datasets compared to randomized data (grey) that simulates the absence of coordination between neighboring genomic bins. **j,** Spearman correlation between principal component (PC) 1 of Hi-C contact matrices and PC1 (red) and PC2 (blue) calculated from LAD coordination matrices.

**Extended Data Figure 3.**
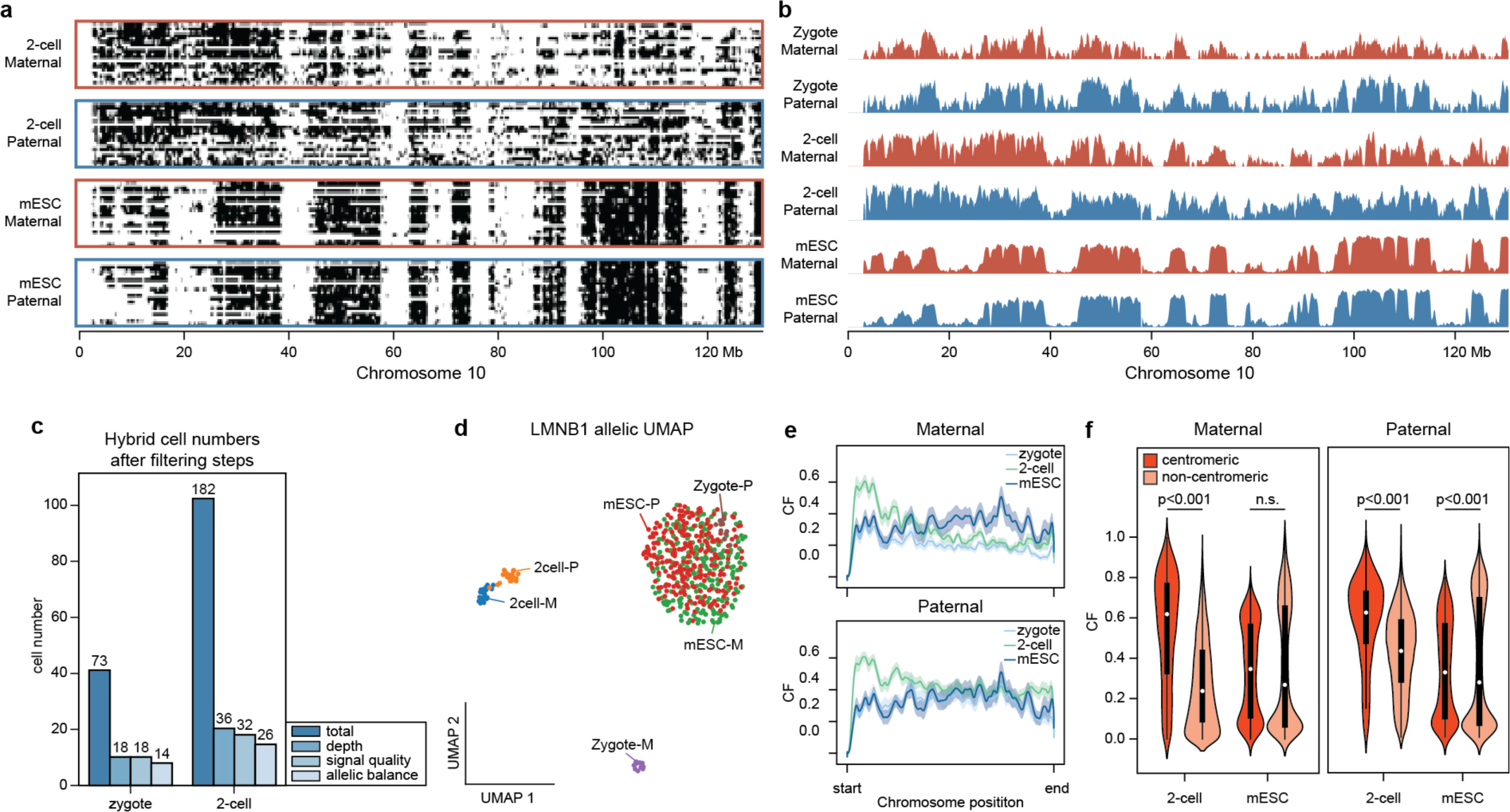
Characterization of single-cell LAD profiles split by parental allele. **a,** Heatmap of binarized single-cell LAD profiles separated into maternal (red box) and paternal (blue box) allele for the 2-cell stage and mESCs along the entire chromosome 10 and odered by unique number of GATCs (DamID depth). All 2-cell samples that passed quality thresholds are pictured while a selection of the same number of mESC samples with highest DamID depth was made for visualization purposes. **b,** Average LMNB1 CF for the maternal (red) and paternal (blue) alleles per stage along the entire chromosome 10. **c,** Cell numbers obtained from hybrid mouse crosses that pass sequential filtering steps: unique number of GATCs (depth), signal quality and allelic balance where the expectation is to recover a similar number of reads from each allele per single cell. **d,** UMAP based on Dam-LMNB1 single-cell readout separated by allele. **e,** Smoothed mean (1000-Mb Gaussian kernel) of LMNB1 contact frequency (CF) distribution across all autosomal chromosomes scaled to the same size per stage and split by allele (maternal - top and paternal – bottom). **f,** Violin plot depicting CF values separated by allele in the first 30 Mb versus the remaining of the chromosome for the 2-cell stage and mESCs. Two-sided Wilcoxon rank-sum test was performed (centromeric bins, n=570; non-centromeric bins, n=24048; maternal 2-cell p-value<2.225×10^-308^, maternal mESC, p-value=0.59, paternal 2-cell p-value<2.225×10^-308^, paternal mESC p-value=0.00024).

**Extended Data Figure 4.**
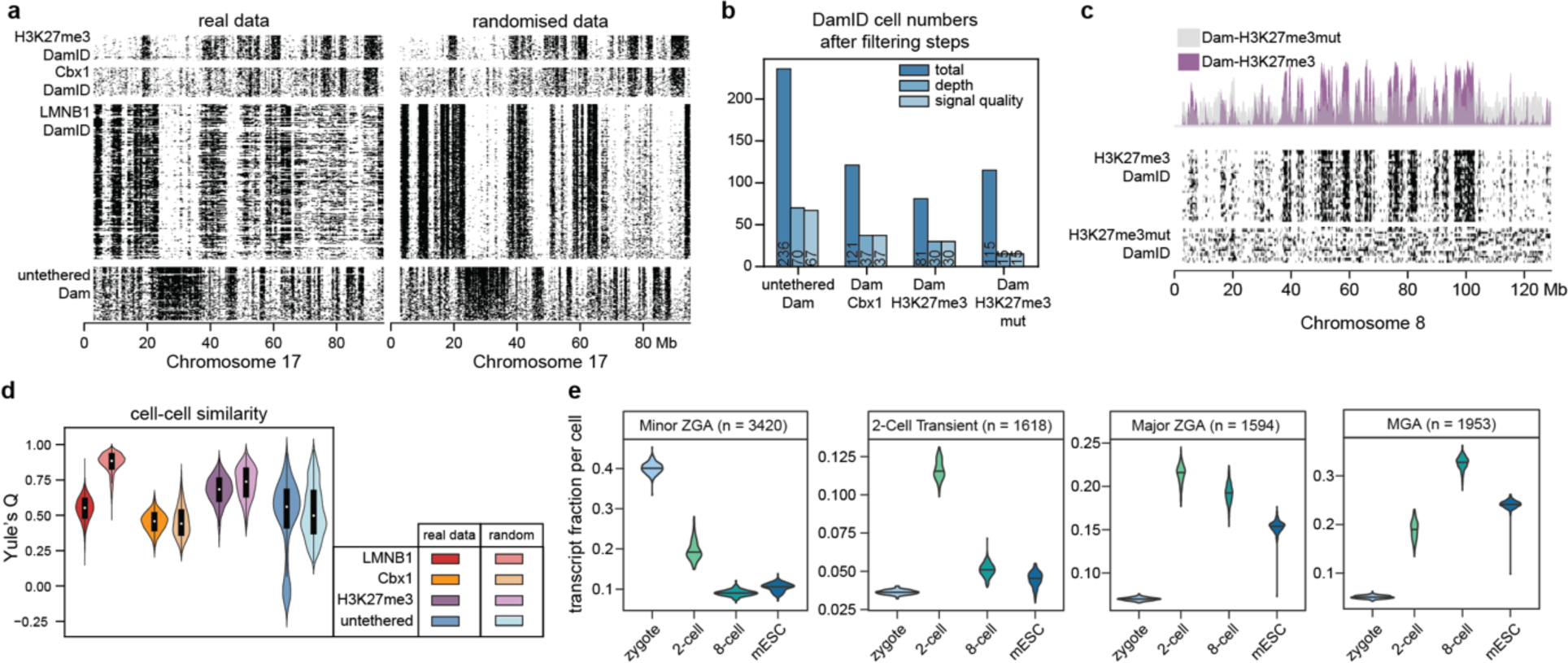
Validation and analysis of single-cell epigenetic marks at the 2-cell stage. **a,** Heatmap of binarized single-cell profiles for Dam-LMNB1, Dam-H3K27me3, Dam-Cbx1 across the entire chromosome 17 ordered by decreasing unique number of GATCs. All cells that passed quality control thresholds are plotted with the size of the heatmap corresponding to relative sample size between constructs (left side). Corresponding randomized data is plotted on the right-side randomized data using the same strategy as in Extended Data Figure 2h. This data is used to correct cell-cell similarity calculations for construct-specific noise and sparsity levels. **b,** Cell numbers that pass sequential filtering steps: unique number of GATCs and signal quality per construct. **c,** Comparison between DamID performed with the Dam-H3K27me3 construct or the mutant version that should not bind H3K27me3. Both binarized single-cell heatmaps (bottom) and corresponding CFs (top) are shown along the entire chromosome 8. **d,** Violin Plot showing Yule’s Q for real data and randomized data (both depicted in a) per construct. Two-sided Wilcoxon rank-sum test was performed (LMNB1, n=196; Cbx1, n=37; H3K27me3, n=30; untethered, n=67) (Extended Data Table 4). **e,** Transcript fraction per cell of gene categories defined by their expression dynamics during early development as defined by Park et al. (2013)^4^^4^ and grouped by stage. ZGA, zygotic genome activation; MGA, mid-preimplantation gene activation.

**Extended Data Figure 5.**
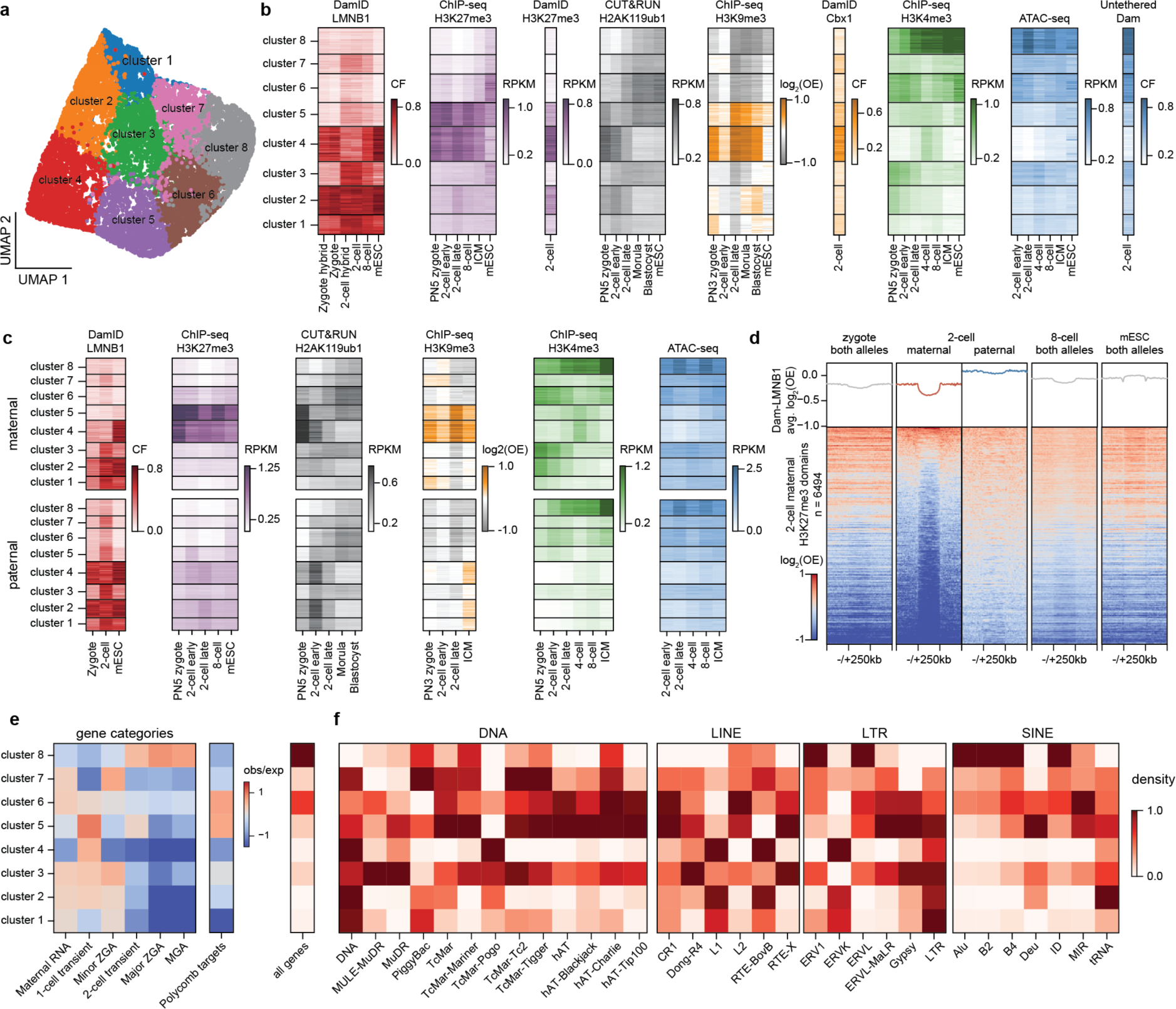
Characterization of genomic regions based on genome-NL association and chromatin marks. **a,** UMAP of genomic bins colored by cluster. **b,** Heatmap with LMNB1, H3K27me3, H2Aub, H3K9me3, H3K4me3 and ATAC-seq values per genomic bin grouped by genomic cluster. **c,** same as b split by allele: maternal on the top and paternal on the bottom. The scale of each measurement is the same for the two alleles. **d,** Enrichment of LMNB1 signal over H3K27me3 maternal domains called from H3K27me3 2-cell stage ChIP-seq data and surrounding upstream and downstream 250 kb. Heatmaps at the bottom show LMNB1 enrichment per domain and line plots on top show average enrichment. **e,** Heatmap showing observed/expected values of different gene categories described in Park et al (2013)^44^, Polycomb targets^45^ and coding gene density for each cluster. **f,** Heatmap showing repeat density of each genomic cluster split by repeat family.

**Extended Data Figure 6.**
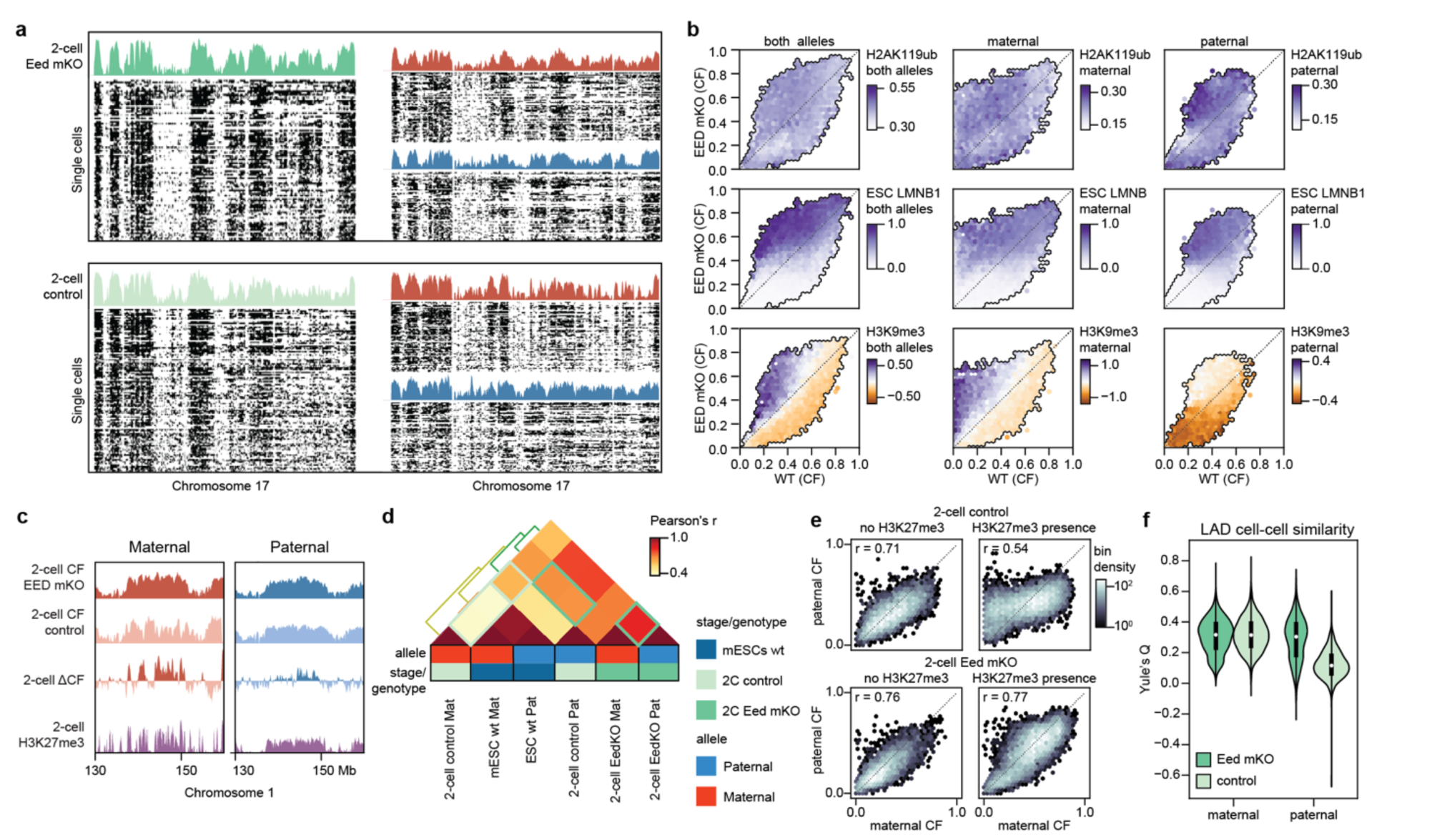
Effect of *Eed* maternal KO on nuclear lamina association at the 2-cell stage. **a,** Single-cell heatmaps of binarized LMNB1 profiles of 2-cell *Eed* mKO (top) or control (bottom) with corresponding CF values per condition along chromosome 17 of both alleles (left) or separated alleles (right). **b,** Correspondence between LMNB1 values in the *Eed*-mKO condition and the control condition for both alleles (left), or maternal and paternal alleles separately (right). The color scale refers to the corresponding combined or allele-specific H2AK119ub1, mESC LMNB1 or H3K9me3 values. **c,** Example genomic region where genome-NL association changes upon *Eed* mKO. LMNB1 CF is plotted for both *Eed* mKO and control as well as the differential CF and H3K27me3 at the 2-cell stage on the maternal (left) and paternal (right) alleles. **d,** Heatmap of Pearson’s correlation between allele-separated LMNB1 profiles of 2-cell *Eed* mKO, 2-cell control and wt mESCs. The KO condition shows higher correlation 1) between maternal and paternal LAD profiles and 2) with mESC LAD profiles as highlighted by green boxes. **e,** Correspondence between maternal and paternal CF in genomic regions containing H3K27me3 (right) or not (left) in either the control 2-cell condition (top) or in the *Eed* mKO (bottom). Color scale refers to density of genomic bins. **f,** Violin plots with distribution of Yule’s Q values as a measure of LAD cell-cell similarity for *Eed* mKO (dark green) and control (light green) separated by allele.

## Extended Data Tables

**Extended Data Table 1.** RNA concentrations of constructs for zygote injections.

| **construct** | **dilution** | **stage collected** | **induction** | **collection (hours post hCG)** |
| --- | --- | --- | --- | --- |
| **Dam-LMNB1** | 5ng/uL | zygote | no | 29-31h |
| **Dam-LMNB1** | 10ng/uL | 2-cell | no | 52-55h |
| **Dam-LMNB1** | <u>100</u>, 150 or 200ng/uL* | 8-cell | no | 75-78h |
| **Dam-H3K27me3** | 5 or <u>10</u>ng/uL* | 2-cell | no | 52-55h |
| **Dam-H3K27me3mut** | 10ng/uL | 2-cell | no | 52-55h |
| **untethered Dam-ERT2** | 20ng/uL | 2-cell | tamoxifen 20h | 52-55h |
| **Dam-Cbx1-ERT2** | 20ng/uL | 2-cell | tamoxifen 20h | 52-55h |

\*multiple concentrations were the result of optimizations: underlined is the most successful condition.

**Extended Data Table 2.** Filtering conditions for single-cell DamID data.

| **Dam construct** | **Embryonic stage** | **# GATC threshold** | **IC threshold** |
| --- | --- | --- | --- |
| Dam-LMNB1 | Zygote | ≥3,000 | ≥1.4 |
| Dam-LMNB1 | 2-cell embryo | ≥10,000 | ≥1.4 |
| Dam-LMNB1 | 8-cell embryo | ≥10,000 | ≥1.4 |
| Dam-LMNB1 | mESC | ≥10,000 | ≥1.4 |
| Dam-only | 2-cell embryo | ≥10,000 | ≥1.2 |
| Dam-H3K27me3 | 2-cell embryo | ≥10,000 | ≥1.2 |
| Dam-H3K27me3mut | 2-cell embryo | ≥10,000 | ≥1.2 |
| Dam-Cbx1 | 2-cell embryo | ≥5,000 | ≥1.2 |

**Extended Data Table 3.** External dataset accession numbers.

| Accession | Technique |
| --- | --- |
| GEO: GSE56697 | MethylC-seq (5mCpG) |
| GEO: GSE66390 | ATAC-seq |
| GEO: GSE71434 | ChIP-seq - H3K4me3 |
| GEO: GSE73952 | ChIP-seq - H3K4me3 & H3K27me3 |
| GEO: GSE76687 | ChIP-seq - H3K27me3 |
| GEO: GSE82185 | Hi-C |
| GEO: GSE97778 | ChIP-seq - H3K9me3 |
| GEO: GSE112551 | DamID – LMNB1 |
| GEO: GSE153496 | CUT&RUN – H2AK119ub1 |
| ENCODE: ENCSR857MYS | ChIP-seq – H3K9me3 |

**Extended Data Table 4.** Wilcoxon test for original vs randomized data.

| Sample 1 | Sample 2 | Wilco x stat | pval | Mean S1 | Mean S2 | Median S1 | Median S2 |
| --- | --- | --- | --- | --- | --- | --- | --- |
| zygote - original | zygote - randomised | -42.50 | 0* | 0.71 | 0.81 | 0.75 | 0.84 |
| 2cell - original | 2cell - randomised | -167.8 | 0* | 0.54 | 0.87 | 0.55 | 0.88 |
| 8cell - original | 8cell - randomised | -129.2 | 0* | 0.50 | 0.72 | 0.49 | 0.73 |
| mESC - original | mESC - randomised | -202.3 | 0* | 0.79 | 0.93 | 0.81 | 0.93 |
| Dam-LMNB1 - original | Dam-LMNB1 - randomised | -167.8 | 0* | 0.54 | 0.87 | 0.55 | 0.88 |
| Dam-only - original | Dam-only - randomised | 2.76 | 0.0057 | 0.49 | 0.52 | 0.56 | 0.49 |
| Dam-2E12 - original | Dam-2E12 - simulated | -7.20 | 5.80E-13 | 0.67 | 0.72 | 0.68 | 0.74 |
| Dam-CBX1_3 - original | Dam-CBX1_3 - simulated | 1.51 | 0.13 | 0.45 | 0.45 | 0.46 | 0.44 |

\*p-values of 0 indicate a value lower than 2.225e-308

**Extended Data Table 5.**
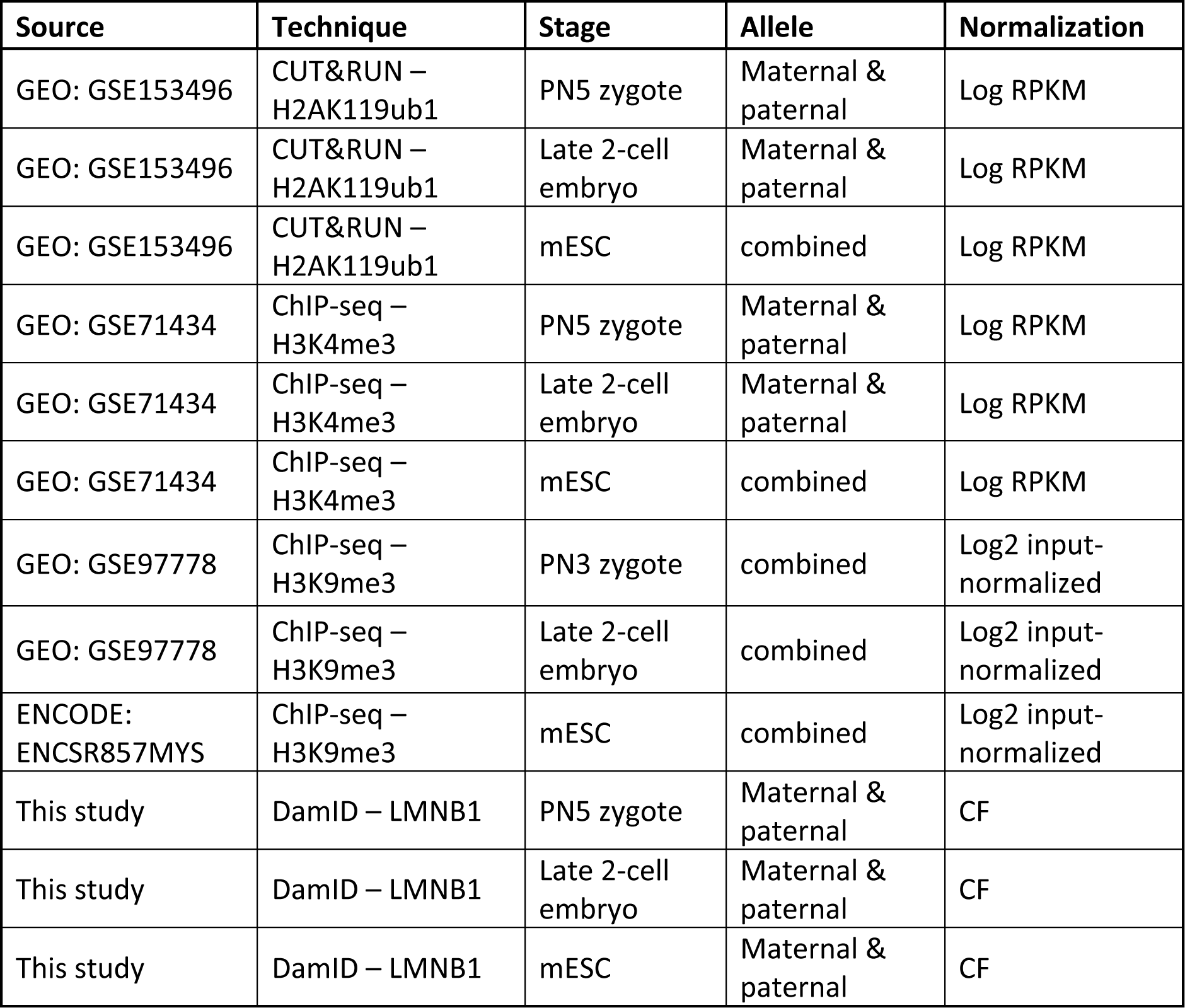
Data used for genomic bin clustering.

| Source | Technique | Stage | Allele | Normalization |
| --- | --- | --- | --- | --- |
| GEO: GSE153496 | CUT&RUN – H2AK119ub1 | PN5 zygote | Maternal & paternal | Log RPKM |
| GEO: GSE153496 | CUT&RUN – H2AK119ub1 | Late 2-cell embryo | Maternal & paternal | Log RPKM |
| GEO: GSE153496 | CUT&RUN – H2AK119ub1 | mESC | combined | Log RPKM |
| GEO: GSE71434 | ChIP-seq – H3K4me3 | PN5 zygote | Maternal & paternal | Log RPKM |
| GEO: GSE71434 | ChIP-seq – H3K4me3 | Late 2-cell embryo | Maternal & paternal | Log RPKM |
| GEO: GSE71434 | ChIP-seq – H3K4me3 | mESC | combined | Log RPKM |
| GEO: GSE97778 | ChIP-seq – H3K9me3 | PN3 zygote | combined | Log2 input-normalized |
| GEO: GSE97778 | ChIP-seq – H3K9me3 | Late 2-cell embryo | combined | Log2 input-normalized |
| ENCODE: ENCSR857MYS | ChIP-seq – H3K9me3 | mESC | combined | Log2 input-normalized |
| This study | DamID – LMNB1 | PN5 zygote | Maternal & paternal | CF |
| This study | DamID – LMNB1 | Late 2-cell embryo | Maternal & paternal | CF |
| This study | DamID – LMNB1 | mESC | Maternal & paternal | CF |

**Extended Data Table 6.**
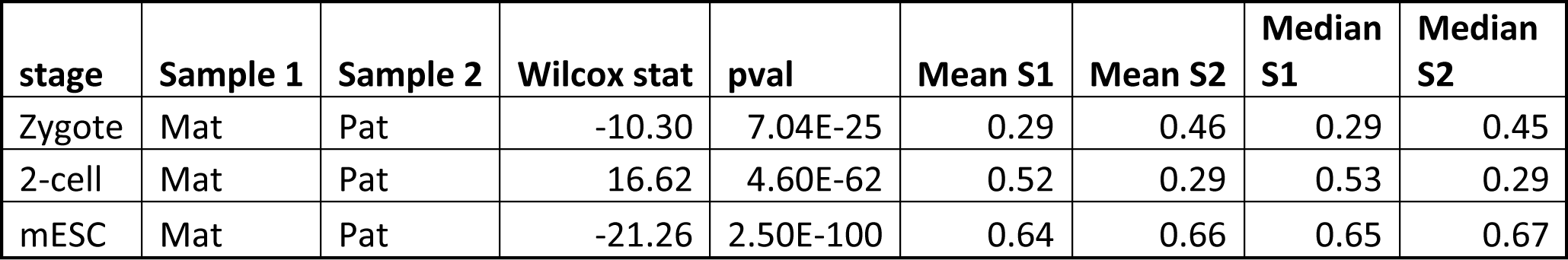
Wilcoxon test for maternal vs paternal cell-cell similarity.

| stage | Sample 1 | Sample 2 | Wilcox stat | pval | Mean S1 | Mean S2 | Median S1 | Median S2 |
| --- | --- | --- | --- | --- | --- | --- | --- | --- |
| Zygote | Mat | Pat | -10.30 | 7.04E-25 | 0.29 | 0.46 | 0.29 | 0.45 |
| 2-cell | Mat | Pat | 16.62 | 4.60E-62 | 0.52 | 0.29 | 0.53 | 0.29 |
| mESC | Mat | Pat | -21.26 | 2.50E-100 | 0.64 | 0.66 | 0.65 | 0.67 |

## Data availability

All genomic and transcriptomic data generated in this study has been deposited at the Gene Expression Omnibus under accession number GSE218598.

## Code Availability

All data analysis code is available upon request.

## Acknowledgements

We would like to thank all the members of the Kind laboratory for their comments throughout the project and their critical reading of the manuscript. We thank Evgeniy A. Ozonov for advice on data analysis. This work was supported by an ERC Starting grant EpiID (ERC Stg EpiID-678423) and ERC Consolidator grant FateID (ERC CoG-101002885) and an NWO-ENW VIDI grant (161.339). The Oncode Institute is partially funded by the KWF Dutch Cancer Society. I.G. was supported by an EMBO Long-Term Fellowship (ALTF1214-2016), Swiss National Science Fund grant (P400PB_186758) and NWO-ENW Veni grant (VI.Veni.202.073). The lab of A.H.M.F.P. has received funding from the Novartis Research Foundation and the European Research Council (ERC) under the European Union’s Horizon 2020 research and innovation programme (grant agreement ERC-AdG 695288 -Totipotency). In addition, we would like to thank the Hubrecht Sorting Facility as well as the Utrecht Sequencing Facility (USEQ), subsidized by the University Medical Center Utrecht.

## Author contributions

I.G., F.J.R. and J.K. designed the study. F.J.R. performed all data analysis with input from I.G. All embryo and scDam&T-seq experiments were performed by I.G. unless otherwise stated with assistance from F.C.G. and R.E.v.B.. Y.K.K. performed the embryo experiments for the *Eed* mKO line and corresponding control with supervision from A.H.M.F.P.. S.J.A.L. and E.B. generated the mESC line expressing Dam-LMNB1 and performed the mESC scDam&T-seq experiment. I.G. wrote the first draft of the manuscript with editing by J.K. All authors reviewed and edited the manuscript.

## Competing interest declaration

The authors declare no competing interests.

Correspondence and material requests should be addressed to Jop Kind or Isabel Guerreiro.

